# Chronic THC vapor inhalation rescues hyperalgesia in rats with chronic inflammation and produces sex-specific alterations in midbrain neuronal activity

**DOI:** 10.1101/2021.09.23.461562

**Authors:** Leslie K. Kelley, Savannah H.M. Lightfoot, Matthew N. Hill, Nicholas W. Gilpin, Jason W. Middleton

**Affiliations:** Departments of Physiology, and Health Sciences Center, New Orleans, LA 70112; Alcohol and Drug of Abuse Center of Excellence, LSUHSC, New Orleans, LA; Hotchkiss Brain Institute, Departments of Cell Biology and Anatomy & Psychiatry, University of Calgary, Calgary, AB, Canada T2N 4N1; Neuroscience Center of Excellence, Louisiana State University Health Science Center, New Orleans, LA 70112; Southeast Louisiana VA Healthcare System, New Orleans, LA; Departments of Cell Biology and Anatomy, Louisiana State University and Health Sciences Center, New Orleans, LA 70112

## Abstract

In an effort to reduce reliance on opioids for the treatment of pain in the clinic, ongoing work is testing the utility of cannabinoid drugs as a potential alternative for treatment of chronic pain. We tested chronic THC vapor inhalation effects on thermal nociception and mechanical sensitivity, as well as midbrain (i.e., ventrolateral periaqueductal gray [vlPAG]) neuronal function, in adult male and female Wistar rats with chronic inflammatory pain (CFA treatment). We report that chronic THC vapor inhalation rescues both thermal hyperalgesia and mechanical hypersensitivity in males treated with CFA, but only thermal hyperalgesia in CFA females. Most of the anti-hyperalgesic effects of chronic THC vapor were still observable 24 hours after cessation of the last THC exposure. We also report that chronic THC vapor inhalation modulates intrinsic and synaptic properties of vlPAG neurons, including reductions in action potential firing rate and spontaneous inhibitory synaptic transmission in males, and that these effects occur specifically in neurons that respond to current input with a “delayed” firing phenotype. Treatment with CFA led to increased firing rate and increased sIPSC amplitude in vlPAG neurons of female rats, and chronic THC vapor rescued sIPSC amplitudes to control levels – these effects were specific to vlPAG neurons categorized as having an “onset” firing phenotype. Ongoing work is exploring sex-specific mechanisms (e.g., CB1 receptor) of THC vapor rescue effects in the vlPAG of rats treated with CFA, and further exploring the vlPAG cell types impacted by CFA treatment and chronic THC vapor inhalation.

**Significance Statement:** Many adults in the U.S. with pain self-medicate with THC and cannabis, and many humans use e-cigarette type devices filled with cannabis extracts to self-administer THC and other constituents of the marijuana plant. Until recently, most rodent studies of THC effects on brain and behavior have used injection procedures and male rats. Here, we tested the effect of chronic THC vapor inhalation on pain-related behaviors and midbrain neural circuit function in adult male and female Wistar rats. As predicted, chronic THC vapor inhalation rescued chronic inflammatory pain effects on behavior and midbrain neuronal function.

## Introduction

Cannabinoids are a promising potential alternative to opioids for the treatment of chronic pain (Nielsen et al., 2017). Interestingly, chronic pain is the most common reason reported for using medicinal cannabis (Baron et al., 2018). With medicinal cannabis approved in thirty-eight U.S. states and recreational cannabis approved in twenty-one U.S. states as of 2023 many adults with pain self-medicate with THC and cannabis (Reiman et al., 2017; Sturgeon et al., 2020), and many cannabis users are increasingly using e-cigarette type devices filled with cannabis extracts to self-administer THC and other constituents of the marijuana plant (Morean et al., 2015; Mammen et al., 2016; Morean et al., 2017). Evidence collected in humans suggests that cannabis is analgesic and therefore may hold promise as a therapeutic substitute for opioids in the clinic (Boehnke et al., 2016; Bradford and Bradford, 2017; Cooper et al., 2018). Despite the prevalent use of THC-containing products for treatment of pain, there is a huge knowledge gap regarding the long-term safety, efficacy, indications, off-target effects and precise mechanisms and sites of action for vaporized THC as a therapeutic.

One promising strategy for reducing pain is via modulation of descending pain modulation circuits (Millan, 2002; Tracey and Mantyh, 2007). One potential point of intervention is at the level of the ventrolateral periaqueductal gray (vlPAG), which receives ascending pain information and modulates incoming pain via descending projections. Chronic inflammatory pain decreases tonic inhibitory current and increases spontaneous inhibitory postsynaptic current frequency in vlPAG neurons in female rats (Tonsfeldt et al., 2016) and increases action potential firing in phasic-firing vlPAG neurons in male and female rats (McPherson et al., 2021). Opioids reduce pain, in part, via activation of descending vlPAG outputs to the rostral ventrolateral medulla (RVM) in adult male rats (Basbaum and Fields, 1984; Hirakawa et al., 1999). Activity of the vlPAG is upregulated by opioid receptor agonists in adult male rats (Loyd et al., 2007), and activation of mu-opioid receptors disinhibits vlPAG neurons through local GABAergic inputs in adult male mice (Li et al., 2016).

While the mechanisms underlying the anti-nociceptive effects of cannabinoids are not completely understood, it is known that cannabinoids reduce inflammation and modulate pain via actions in central pain pathways (Guindon and Hohmann, 2009; Woodhams et al., 2017). More specifically, cannabinoids are thought to reduce pain via activation of cannabinoid type-1 receptors (CB1Rs) that are found in many of the same brain regions as opioid receptors (Befort, 2015), including in the vlPAG where they are highly expressed (Wilson-Poe et al., 2012). Furthermore, infusions of synthetic cannabinoid receptor agonists into the ventricles, the PAG and the RVM all produce analgesia in naïve male rats (Lichtman et al., 1996; Meng et al., 1998). It is not known how THC vapor inhalation alters vlPAG function in experimentally naïve animals or in animal models of chronic inflammatory pain.

Many humans vape exogenous cannabinoid compounds (Giroud et al., 2015; Lee et al., 2016; Trivers et al., 2019), but animal studies have mainly relied on intraperitoneal injections of cannabinoids in males to study drug effects on pain-related (and other) outcomes (Britch and Craft 2021). That said, cannabinoid vapor inhalation is being increasingly tested for its effects on nociception, body temperature and drug discrimination in rats (Hložek et al., 2017; Nguyen et al., 2018; Moore and Weerts, 2021; Moore et al., 2021; Wiley et al., 2021). It is clear that route of drug administration and sex influence the pharmacokinetic and behavioral effects of cannabis in humans and animals (Sholler et al., 2022, 2022; Baglot et al., 2021; Zamarripa et al., 2022). Many prior studies have tested the effect of exogenous cannabinoids on acute rather than chronic pain (Cooper and Craft 2018) and much of this work has been performed in male animals.

One study reported that, relative to male rats treated with CFA, systemic and intra-plantar THC injection more effectively reduces thermal hyperalgesia and/or mechanical allodynia in CFA-treated female Sprague-Dawley rats, whereas both routes of THC injection more effectively reduce paw edema in male rats (Craft et al., 2013). Another study reported that systemically injected THC dose-dependently reduces pain-related behaviors in CFA-treated male and female rats but does not reduce hindpaw edema, with little evidence for sex differences or tolerance to those effects after repeated treatment (Britch et al., 2020), and similar effects were observed in a more recent study by the same group (Britch and Craft, 2023). Finally, one very recent study reported that vaporized THC-dominant cannabis extract reduces mechanical allodynia and paw edema similarly in male and female Sprague-Dawley rats treated with CFA, and that tolerance to these effects develop with chronic exposure (Craft et al., 2023). To our knowledge, no studies have examined the effect of vaporized THC on pain-related behavior in CFA-treated rats nor on neuronal function in midbrain regions important for descending pain modulation.

Here, we aimed to test the effect of chronic THC vapor inhalation on chronic inflammatory pain-related outcomes and vlPAG neuronal function in male and female rats. We performed all experiments in male and female rats, but because male and female experiments were not performed at the same time, we did not perform direct statistical comparisons of male and female behavioral and neuronal data (but we did include sex as a factor in analyses of THC and metabolite plasma levels). We hypothesized that chronic THC vapor inhalation would have anti-hyperalgesic and anti-allodynic effects in rats with chronic inflammatory pain (i.e., treated with CFA), and that it would produce disinhibition of vlPAG neurons.

## Methods

### Animals

Adult male (N=94) and female (N=66) specific pathogen free Wistar rats (Charles River, Raleigh, NC) were housed in a humidity- and temperature-controlled (22 °C) vivarium on a 12 h light/12 h dark cycle (lights off at 8 a.m.). All rats were housed in groups of two. Rats were 8 weeks old upon arrival and acclimated for one week before start of experiments. Rats were handled daily prior to initiation of experimental protocols. Behavioral testing and vapor exposure occurred during the dark period. All procedures were approved by the Institutional Animal Care and Use Committee of the Louisiana State University Health Sciences Center and were in accordance with the National Institute of Health guidelines.

### Drugs

Δ^9^-tetrahydrocannabinol (THC) was supplied in ethanolic solution (200mg/mL) by the Research Triangle Institute (RTI) in conjunction with the U.S. National Institute on Drug Abuse (NIDA) Drug Supply Program. THC was administered at a dose of 100 mg THC diluted in propylene glycol (PG) vehicle (1 ml) via vapor inhalation during 1-hour daily sessions that occurred over 10 consecutive days.

### Complete Freund’s Adjuvant (CFA)

A localized chronic inflammatory pain state was established by injecting 150 μL of 50% complete Freund’s adjuvant (Sigma) in 0.9% saline subcutaneously into the intraplantar surface of the left hindpaw. This injection produced visible inflammation that resulted in thermal hyperalgesia and mechanical hypersensitivity (see *Results*). Control rats were injected with 150 μL of 0.9% saline into the left hindpaw that produced no inflammation.

### Nociception assays

For Experiment 1, pre-CFA and post-CFA thermal nociception and mechanical sensitivity were assessed over several sessions. The 3 sessions preceding CFA injections were averaged together to obtain pre-CFA baseline thermal nociception and mechanical sensitivity scores for each rat, and those data were used to counterbalance baseline pain-like behaviors in rats assigned to CFA and Saline groups. Following injection of CFA or Saline, rats were tested for 3 additional sessions of thermal nociception and mechanical sensitivity, and the scores obtained in those tests were used to assign rats to one of four groups that were counterbalanced for post-CFA pain-like behaviors. There were four groups of male rats treated with either CFA + 100 mg THC vapor (N=8), Saline + 100 mg THC vapor (N=8), CFA + Vehicle (PG) vapor (N=12), or Saline + Vehicle (PG) vapor (N=12). There were also four groups of female rats treated with either CFA + 100 mg THC vapor (N=10), Saline + 100 mg THC vapor (N=10), CFA + Vehicle (PG) vapor (N=8), or Saline + Vehicle (PG) vapor (N=8).

During the 10 days of THC or PG vapor exposure (1 hour per day), rats were repeatedly tested for either thermal nociception or mechanical sensitivity in the Hargreaves and Von Frey tests respectively, across alternating days, as follows: Hargreaves tests were conducted on vapor days 2,5, and 8; Von Frey tests were conducted on vapor days 3, 6, and 9. All of those tests occurred 5 minutes after the end of vapor inhalation sessions. Hargreaves and Von Frey tests were repeated one final time on day 11 of the protocol at a time point that coincided with 24 hours after the 10^th^ and final vapor exposure session.

### Hargreaves thermal sensitivity testing

Before all pre-CFA/saline and post-CFA/saline baseline nociception testing, animals were habituated to the behavior room for 30 minutes. Once room habituation was completed, rats were placed in 4 × 8 × 5 inch clear plexiglass enclosures on top of a glass pane suspended 8 inches above the table top. Thermal nociception threshold was measured by Hargreaves plantar test (Hargreaves Apparatus, Model 309, IITC Life Sciences), which measures the latency for each rat to withdraw its hindpaw when stimulated by a halogen lamp heat source from underneath a glass pane. Paw withdrawal latency (PWL) in response to the application of thermal heat was recorded in seconds with a cutoff time of 20 seconds to prevent potential burn. Four readings were obtained from each rat in one session, two from each hindpaw in an alternating manner with at least 1 min between the readings. The readings were then averaged to determine a thermal nociception score for each hindpaw from each rat.

### Von Frey mechanical sensitivity testing

Mechanical sensitivity was determined using automated Von Frey equipment (Ugo Basile, Italy) and obtaining pre-CFA and post-CFA baseline hindpaw withdrawal thresholds. On experimental days where Von Frey and Hargreaves testing occurred on the same day (pre-CFA and post-CFA baseline days and day 11 post-vapor), Von Frey testing always occurred following Hargreaves testing. On these days, rats were acclimated to the test room for 30 minutes, tested for thermal nociception in the Hargreaves test, then acclimated for 15 minutes to individual plexiglass compartments set on top of a mesh stand and tested for mechanical sensitivity in the Von Frey test. In the Von Frey test, an automated device is used that applies a steel filament to the plantar surface of the hindpaw with gradually increasing force until the hindpaw is withdrawn (the withdrawal force is rounded to the nearest 0.1 g). For each rat on each test day, the filament was applied to each hindpaw twice (four total applications) in an alternating manner with at least 1 minute between each application and measurement. Measured values were averaged to determine a mechanical sensitivity score for each hindpaw from each rat on each test day.

### THC Vapor Delivery

Seven days after the injection of CFA or saline, rats were placed in sealed exposure chambers (35 x 28 x 26 cm) manufactured by LJARI for 1-hour long vapor sessions. An e-vape controller (Model SVS-200; 30.0 watts; La Jolla Alcohol Research, Inc., La Jolla, CA, USA) was scheduled to deliver 6 s puffs every 5 min from a Smok Baby Beast Brother TFV8 sub-ohm tank (fitted with the V8 X-Baby M2 0.25 Ω coil). The volume of PG vehicle or THC 100mg solution used was ∼1 mL per 1-hour session. The chamber air was vacuum-controlled to flow ambient air through an intake valve at ∼0.6 L per minute. The vapor cloud cleared from the chamber by 2 min after the puff delivery. Animals were exposed to THC or vehicle vapor for one hour daily for 10 days. Plasma was collected for analysis via tail bleeds immediately after the 1-hour session ended on days 1 and 10 in Experiment 1.

### Plasma THC analysis

Tail blood samples (∼500 μL) were collected by making a cut 1 mm at the tip of the tail; the same cut was used to collect blood from each rat immediately after the end of the 1-hour vapor session on days 1 and 10. Blood was collected in Eppendorf tubes and kept on ice until centrifugation. After centrifugation, plasma (∼50 μL) was stored at -80°C until analysis.

Full method of THC analysis can be found in (Baglot et al., 2021); in brief, plasma samples were thawed and 100 μL of sample was added directly to borosilicate glass tubes containing 2 ml of acetonitrile and 100 µL of internal standard (IS) stock solutions (IS contain THC-d3, 11-OH-THC-d3 and THC-COOH-d3 were dissolved in acetonitrile at 0.1 mg/mL; all d3 standards from Cerilliant, Round Rock, TX, USA). All samples were then sonicated in an ice bath for 30 min before being stored overnight at -20°C to precipitate proteins. The next day samples were centrifuged at 1800 rpm at 4°C for 3-4 min to remove particulates and the supernatant from each sample was transferred to a new glass tube. Tubes were then placed under nitrogen gas to evaporate. Following evaporation, the tube sidewalls were washed with 250 µL acetonitrile in order to recollect any adhering lipids and then again placed under nitrogen gas to evaporate. Following complete evaporation, the samples were re-suspended in 100 µL of 1:1 methanol and deionized water. Resuspended samples went through two rounds of centrifugation (15000 rpm at 4°C for 20 min) to remove particulates and the supernatant transferred to a glass vial with a glass insert. Samples were then stored at -80°C until analysis by LC-MS / Multiple Reaction Monitoring (MRM). LC-MS/MS analysis was performed using an Eksigent Micro LC200 coupled to an AB Sciex QTRAP 5500 mass spectrometry (AB Sciex, Ontario, Canada) at the Southern Alberta Mass Spectrometry (SAMS) facility located at the University of Calgary, as previously described (Baglot et al., 2021). Analyte concentration (in pmol/µL) were normalized to sample volume and converted to ng/mL.

### Electrophysiology

Under isoflurane anesthesia, rats were transcardially perfused with 100 mL room temperature (∼25°C) NMDG artificial cerebrospinal fluid (aCSF) containing the following (in mM): 92 NMDG, 2.5 KCl, 1.25 NaH_2_PO_4_, 30 NaHCO_3_, 20 HEPES, 25 glucose, 2 thiourea, 0.5 CaCl_2_, 10 MgSO_4_·7 H_2_O, 5 Na-ascorbate, 3 Na-pyruvate. 300 µm-thick coronal sections containing the left vlPAG were collected using a vibratome (Leica VT1200S, Nussloch, Germany). Sections were incubated in NMDG aCSF at 37°C for 12 min, then transferred to a room temperature holding aCSF solution containing the following (in mM): 92 NaCl, 2.5 KCl, 1.25 NaH_2_PO_4_, 30 NaHCO_3_, 20 HEPES, 25 glucose, 2 thiourea, 2 CaCl_2_, 2 MgSO_4_·7 H_2_O, 5 Na-ascorbate, 3 Na-pyruvate. Slices were allowed to recover for one hour prior to recording. Slices were visualized with oblique infrared light illumination, a w60 water immersion objective (LUMPLFLN60X/W, Olympus, Tokyo, Japan) and a CCD camera (Retiga 2000R, QImaging, Surrey, BC, Canada). Data were collected using the acquisition control software package Ephus (Suter et al., 2010).

At the time of recording slices were transferred to slice chamber in the electrophysiology rig in a recording aCSF kept at 30-32°C by an in-line heater. The recording solution consisted of (in mM): 119 NaCl, 2.5 KCl, 1.25 NaH_2_PO_4_, 24 NaHCO_3_, 12.5 glucose, 2 CaCl_2_, 2 MgSO_4_. Whole-cell recordings were performed using borosilicate glass micropipettes (3–7MOhm) filled with internal solution containing (in mM): 130 K-gluconate, 10 HEPES, 10 Na_2_-phosphocreatine, 4 MgCl_2_, 4 Na_2_-ATP, 0.4 Na-GTP, 3 ascorbic acid, 0.2 EGTA (pH 7.25, 290–295 mOsm). Electrical signals were amplified and digitized by a Multiclamp 700B amplifier (Molecular Devices, San Jose, CA). Recordings were sampled at 10 kHz and low pass Bessel filtered with a cutoff of 4 kHz. Signals were further filtered offline in Matlab with a cutoff of 2 kHz.

#### Synaptic Properties

To collect data regarding pre- and postsynaptic alterations resulting from TBI and inhibitor treatment, spontaneous excitatory postsynaptic currents (sEPSCs) were recorded. When whole-cell recording configuration was established, the voltage was clamped at -50 mV and 5 min of spontaneous postsynaptic current was recorded. Blockers of glutamate receptors, CNQX (10 μM) + APV (50 μM) were circulating in the aCSF bath throughout all recordings. Spontaneous inhibitory postsynaptic events were detected in these recordings by thresholding rapid excursions in outward current; the average event amplitude and mean frequency over the 5 min recording period were quantified.

#### Intrinsic Properties

Because intrinsic subthreshold neural properties also contribute to integration of input synaptic signals and neural excitability, injury and/or treatment effects on intrinsic neural properties were assessed; specifically, input resistance, SAG fraction, and resting membrane potential (RMP) were measured. To estimate input resistance, holding current was recorded in voltage clamp mode with membrane voltage clamped at -70 mV. A -5 mV step was applied to the command voltage and the change in holding current, ΔI, was measured. The input resistance was estimated using Ohm’s law, R = ΔV / ΔI. Rebound response to hyperpolarization activated inward current, known as the voltage SAG, was calculated as ΔV2 / ΔV1, where ΔV1 is the difference between pre-step baseline voltage (-70 mV) and the steady state voltage during step current injection, and ΔV2 is the difference between current step steady state voltage and the minimum voltage during current injection. Voltage SAG (Dembrow et al., 2010; Joshi et al., 2015) is reflective of subthreshold current mediated by HCN channels, also known as *I_h_*. RMP was recorded in current clamp mode with injected current set at 0 pA.

#### Action Potential Generation Properties

Properties related to action potential generation were also examined, including spike threshold, firing rate, and firing rate-input gain. The spike threshold is the most depolarized membrane voltage level before the action potential excursion. In this study, it was defined as the voltage at which the second derivative (i.e. curvature) of the voltage was at its local maximal value. Input current steps of varying current amplitudes were delivered and the membrane voltage response to these current step inputs was recorded. Action potentials during each current step were counted and used to construct a firing rate-input (FI) curve. The FI curve gain was quantified by calculating the slope of the curve between 50 and 150 pA.

### Experiment 1: Effect of THC vapor exposure on thermal nociception and mechanical sensitivity

Experiment 1 was conducted in male (N=40) and female (N=38) Wistar rats. Male and female rats were tested separately for thermal nociception and mechanical sensitivity, as described above. Rats were assigned to CFA or Saline groups counterbalanced for baseline nociception scores. Rats were then injected in the left hindpaw with CFA (Males N=20, Females N=19) or saline (Males N=20, Females N=19), tested again for HG and VF, and these scores were used to assign rats to either THC (100 mg) vapor or vehicle (PG) vapor groups. Final treatment groups were as follows: CFA + THC vapor (Males N=8, Females N=10), Saline + THC vapor (Males N=8, N=10), CFA + Vehicle vapor (Males N=12, Females N=9), Saline + Vehicle vapor (Males N=12, Females N=9). Seven days after injection of CFA or saline, rats began 10 days of THC or PG vapor exposure. Rats were exposed to vapor for 1 hour daily, with tail bleeds occurring immediately following the sessions on days 1 and 10. Rats were tested for thermal sensitivity using HG (days 2,5, and 8) or mechanical sensitivity using VF (days 3,6, and 9) five minutes after termination of the vapor session. Rats were once again tested in the HG and VF assays 24 hours after the last vapor exposure session. Data were analyzed using 3-way RM ANOVAs where CFA condition and vapor exposure were between-subjects factors and time was a within-subjects factor: for each outcome measure (e.g., thermal nociception and mechanical sensitivity), one analysis included baseline and three vapor test days to assess acute THC vapor effects on behavior; for each outcome measure, a separate analysis included baseline and the 24-hour post THC vapor time point to assess lasting THC vapor effects on behavior. 2-way RM ANOVAs (vapor condition x time) were also conducted to assess acute and lasting effects in CFA-treated rats only, and again in saline-treated rats only.

### Experiment 2: Effect of THC vapor exposure on plasma THC concentrations

Experiment 2 was conducted using plasma obtained from the rats tested in Experiment 1. Tail blood collections occurred immediately after the end of vapor exposure sessions on days 1 and 10. Plasma samples were frozen and stored at -80°C until MS analysis. Three THC plasma concentration values were identified as outliers using the interquartile range (IQR) test: specifically, they were >1.5 x IQR above the 3^rd^ quartile (Q3) value. For these 3 samples, all measurements (THC and its metabolites) were excluded from analyses. Data were analyzed using 3-way RM ANOVA, where CFA condition and vapor exposure were between-subjects factors and time was a within-subjects factor.

### Experiment 3: Effects of CFA and THC vapor exposure on intrinsic properties and synaptic transmission in vlPAG neurons

Experiment 3 was conducted using a second group of male (N=44) and female (N=28) Wistar rats. Rats were injected in the left hindpaw with either CFA or saline. Seven days after the injection of CFA or Saline rats were exposed to either THC (100 mg) or vehicle (PG) vapor during daily 1-hour sessions for 10 consecutive days. Twenty-four hours after the 10^th^ and final vapor exposure session, rats were sacrificed for electrophysiological brain slice experiments. Analysis of electrophysiological data was performed in Matlab (MathWorks, Natick, MA). The final groups for male rats were as follows: CFA + THC = 55 neurons from 14 rats, CFA + PG Vapor = 41 neurons from 11 rats, Saline + THC = 44 neurons from 8 rats, Saline + PG Vapor = 43 neurons from 11 rats). The final groups for female rats were as follows: CFA + THC = 44 neurons from 10 rats, CFA + PG Vapor = 41 neurons from 10 rats, Saline + THC = 45 neurons from 11 rats, Saline + PG Vapor = 48 neurons from 11 rats).

Electrophysiological data were analyzed using 2-way ANOVAs where CFA condition and vapor exposure were between-subjects factors. Post-hoc pairwise comparisons were performed using Tukey’s honest significant difference (HSD). Statistics were performed on groups where individual samples correspond to individual neurons. The magnitude of main effects (for results with p < 0.05) were quantified using partial eta squared, η_p_^2^, which measures the proportion of variation in the dependent variable associated with its membership in an experimental group (Lakens, 2013).

On average, there were 4.2 neurons per rat across all groups in our data set. Statistical analysis performed on electrophysiological data was done considering each recording parameter as an independent measurement (i.e., N = # of neurons). Neural properties exhibit significant heterogeneity (even within a single cell class), and neural property heterogeneities can be important for determining neural circuit function (Pouille et al, 2009; Padmanabhan and Urban, 2010). One concern, however, is that collecting multiple samples from the same animal may result in pseudo-replication under certain conditions – specifically, when the intra-group variance of individual animal averaged parameters (animal means) is large relative to the individual data point variance within an animal, there is an increased chance of obtaining type I errors (false positives) (Eisner, 2021). One tool to assess this risk is calculating the intraclass correlation coefficient, which quantifies the relative mean discrepancy within groups (Sikkel et al, 2017). In all cases where we observed a significant ANOVA effect, we calculated the ICC for the analyzed electrophysiological parameter and confirmed that significant correlations within a group did not contribute to the significant ANOVA effect in question.

## Results

### Experiment 1

#### CFA produces mechanical hypersensitivity and thermal hyperalgesia in male and female rats

Rats treated with CFA in the left hindpaw exhibited increases in mechanical sensitivity and thermal nociception, as evidenced by lower force thresholds required for (left) hindpaw withdrawal in the von Frey test (**Fig. 2A**, Males: t(38)=11.22, p<0.0001; Females: t(36)=5.73, p<0.0001), and lower (left) hindpaw withdrawal latency times recorded during the Hargreaves test (**Fig. 2B**, Males: t(38)=11.59, p<0.0001, Females: t(36)=5.41, p<0.0001) relative to saline-treated rats. These data are illustrated in the BL timepoint of **Figures 2A and 2B**. CFA-treated rats did not exhibit mechanical hypersensitivity or thermal hyperalgesia in the untreated (right) hindpaw (**Table 1**).

**Figure 1:**
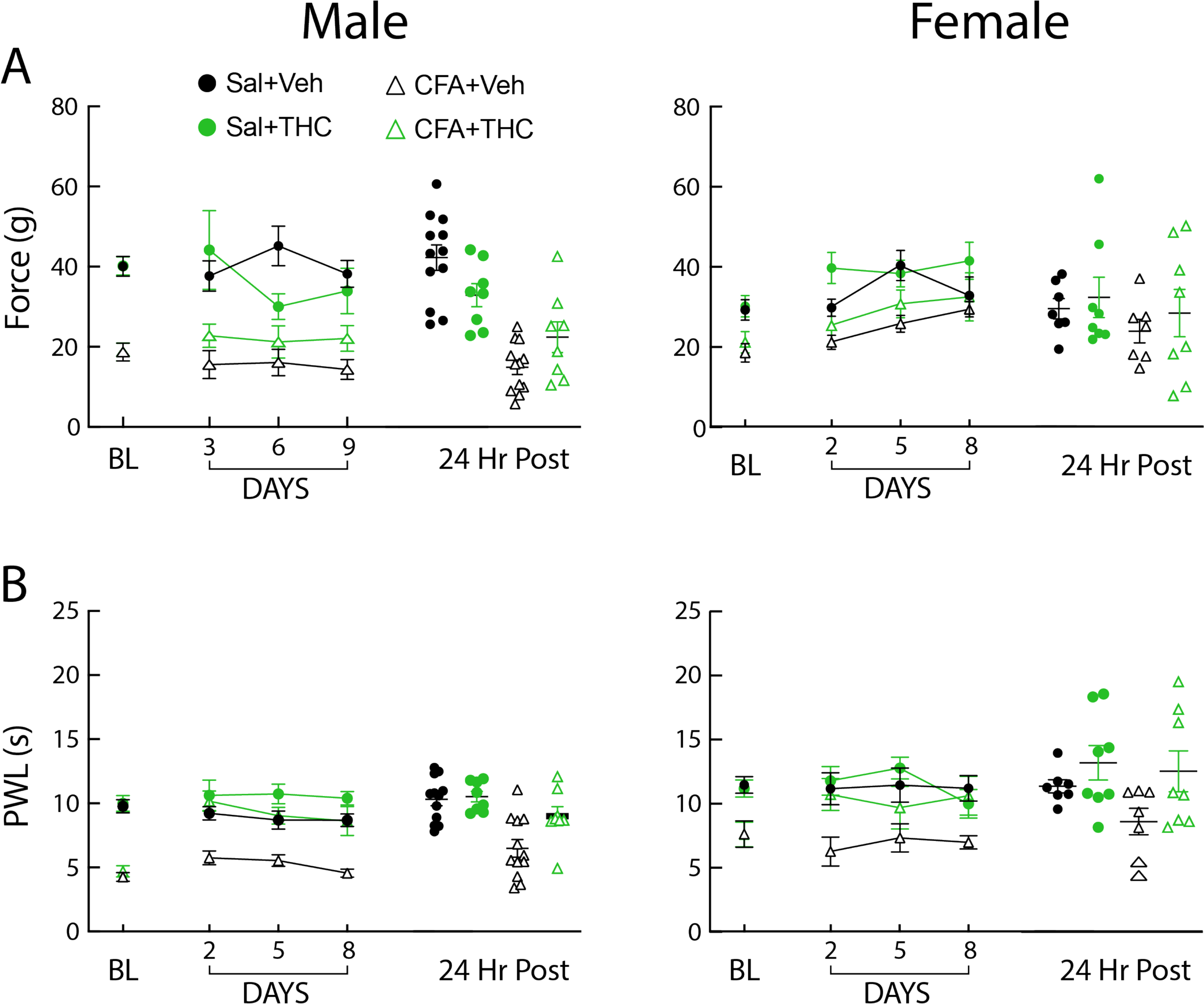
Timelines for Experiment 1 (top) and Experiment 2 (bottom).

**Figure 2:**
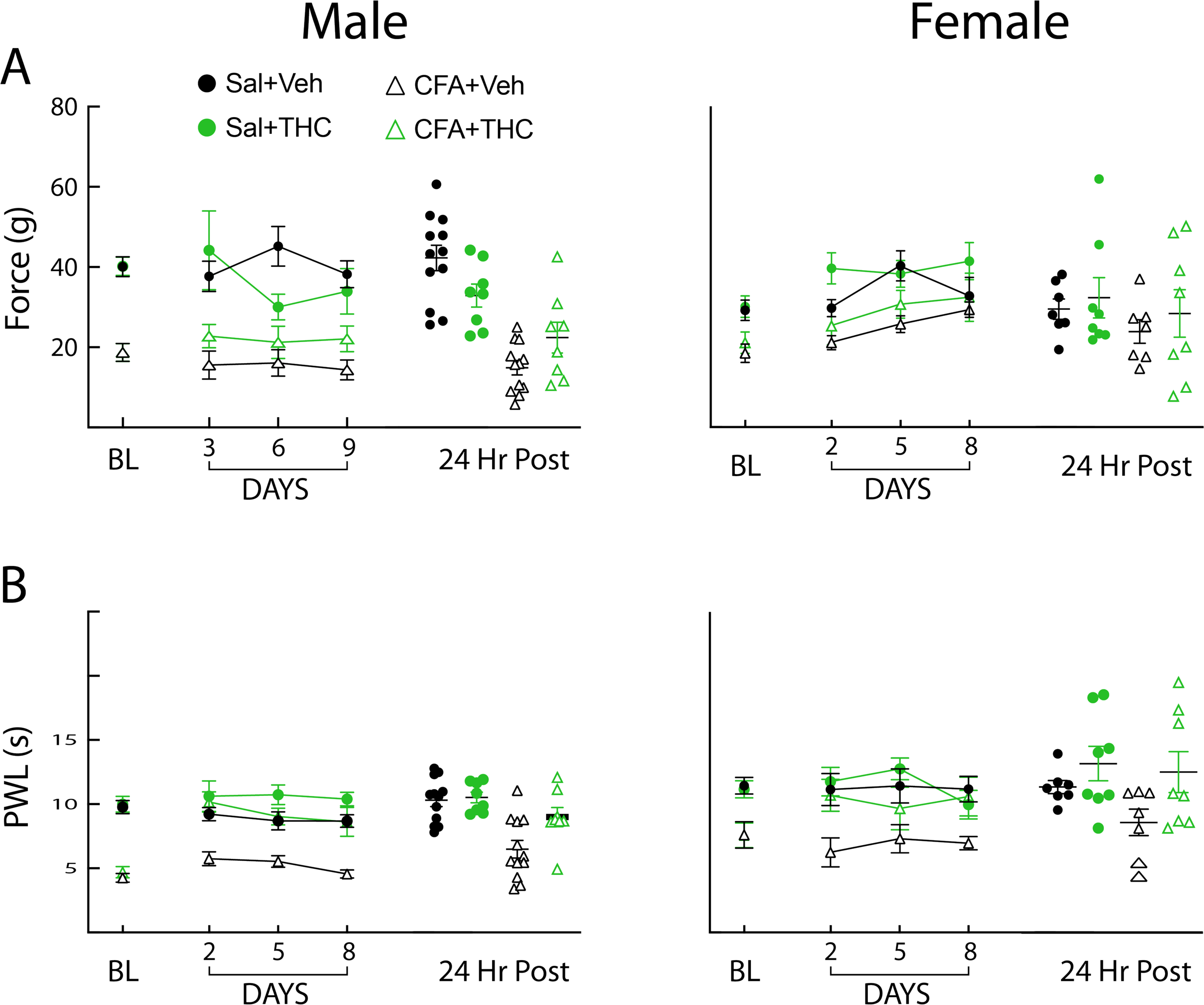
Effects of CFA condition and chronic vapor on mechanical and thermal nociception. (**A**) Behavioral data for Mechanical Sensitivity Baseline, vapor Days 3,6, 9 and 24 hours after the last vapor exposure. 3-Way repeated measures ANOVAs reveal no effect of THC vapor history on mechanical sensitivity. Left: Male rats; Right: Female rats. (**B**) Thermal Nociception Baseline, vapor Days 2,5, 8 and 24 hours after the last vapor exposure. 3-Way repeated measures ANOVAs reveal acute and lasting effects of THC vapor on thermal nociception. Left: Male rats; Right: Female rats.

**Table 1:**
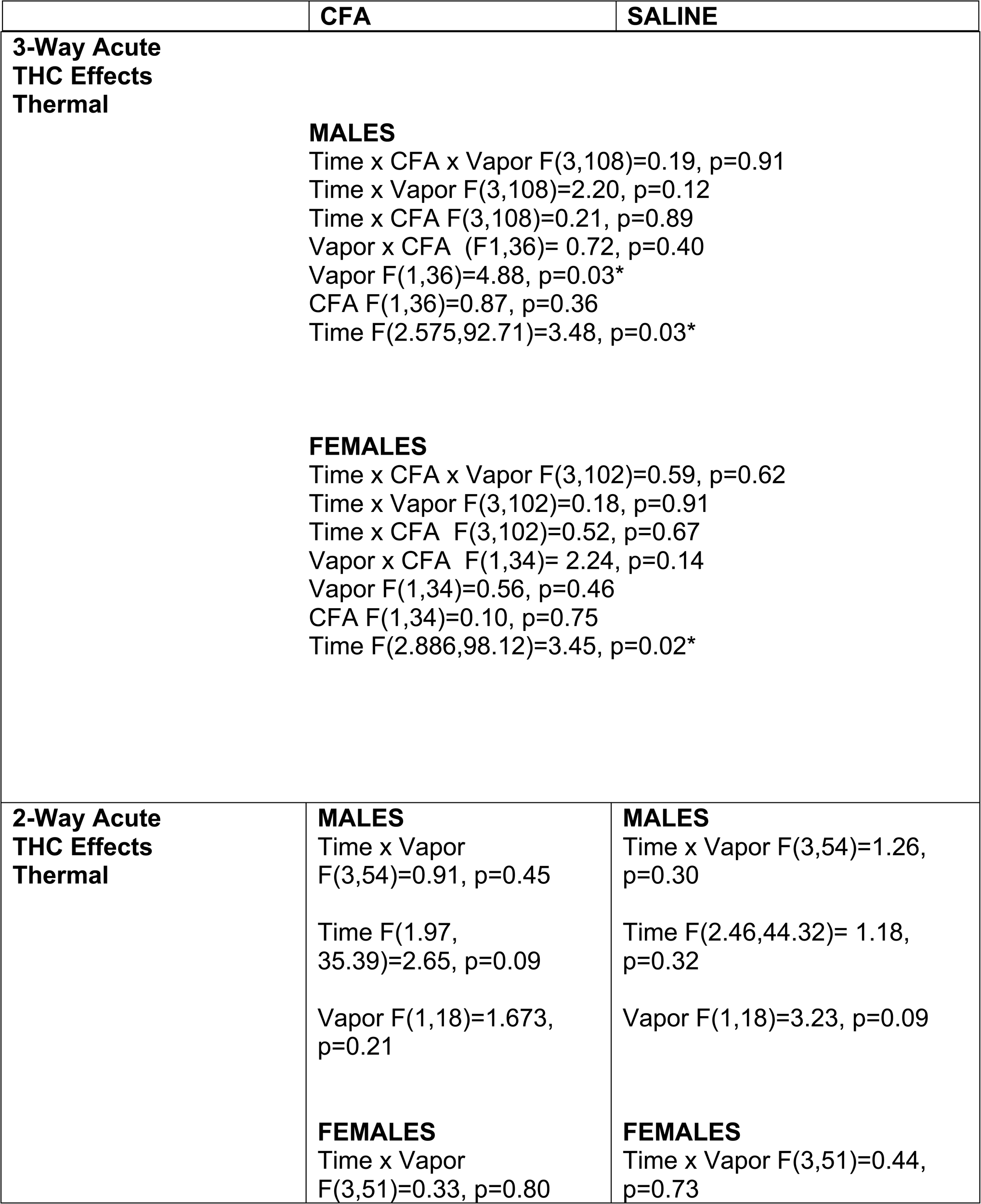
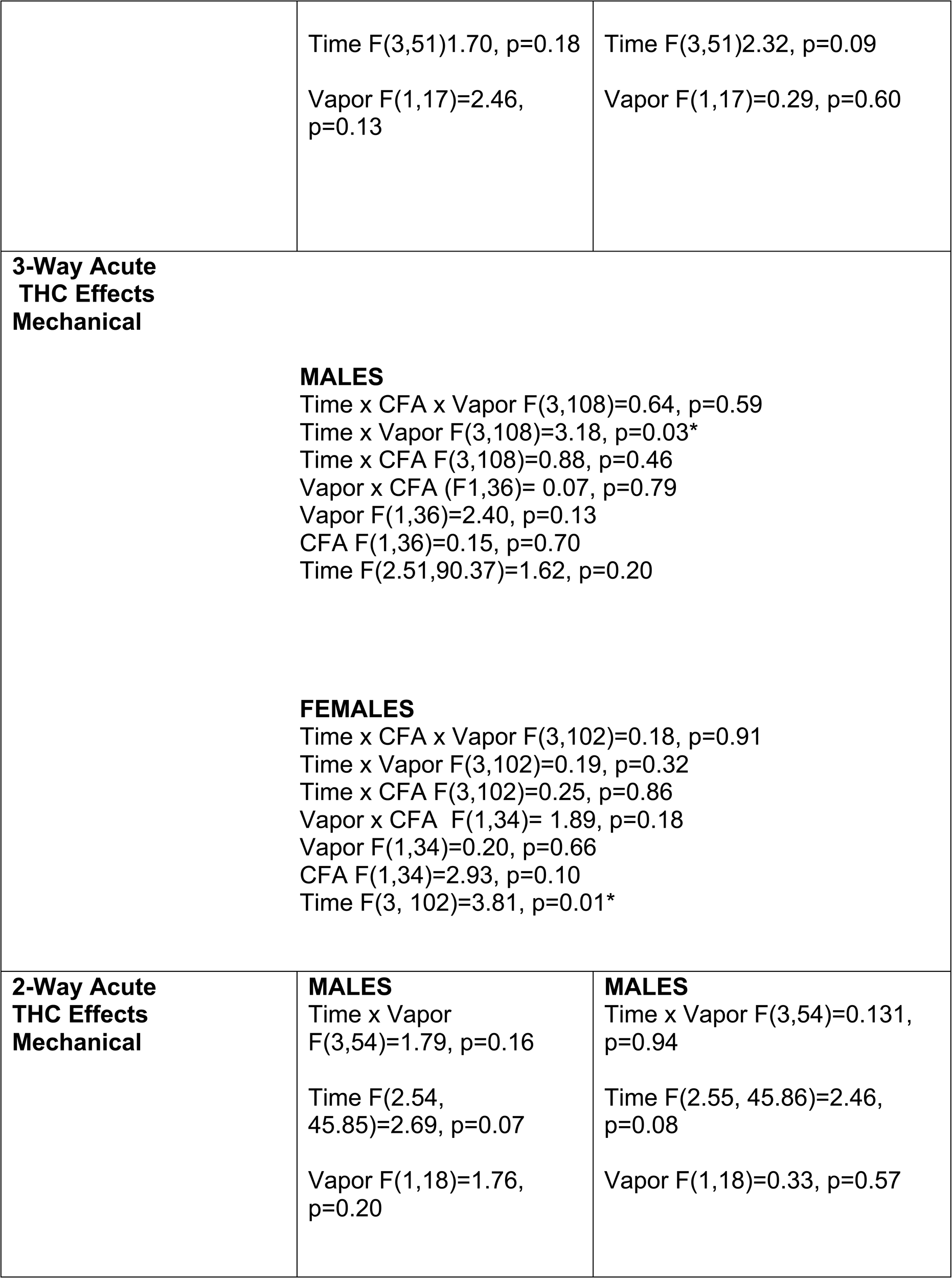
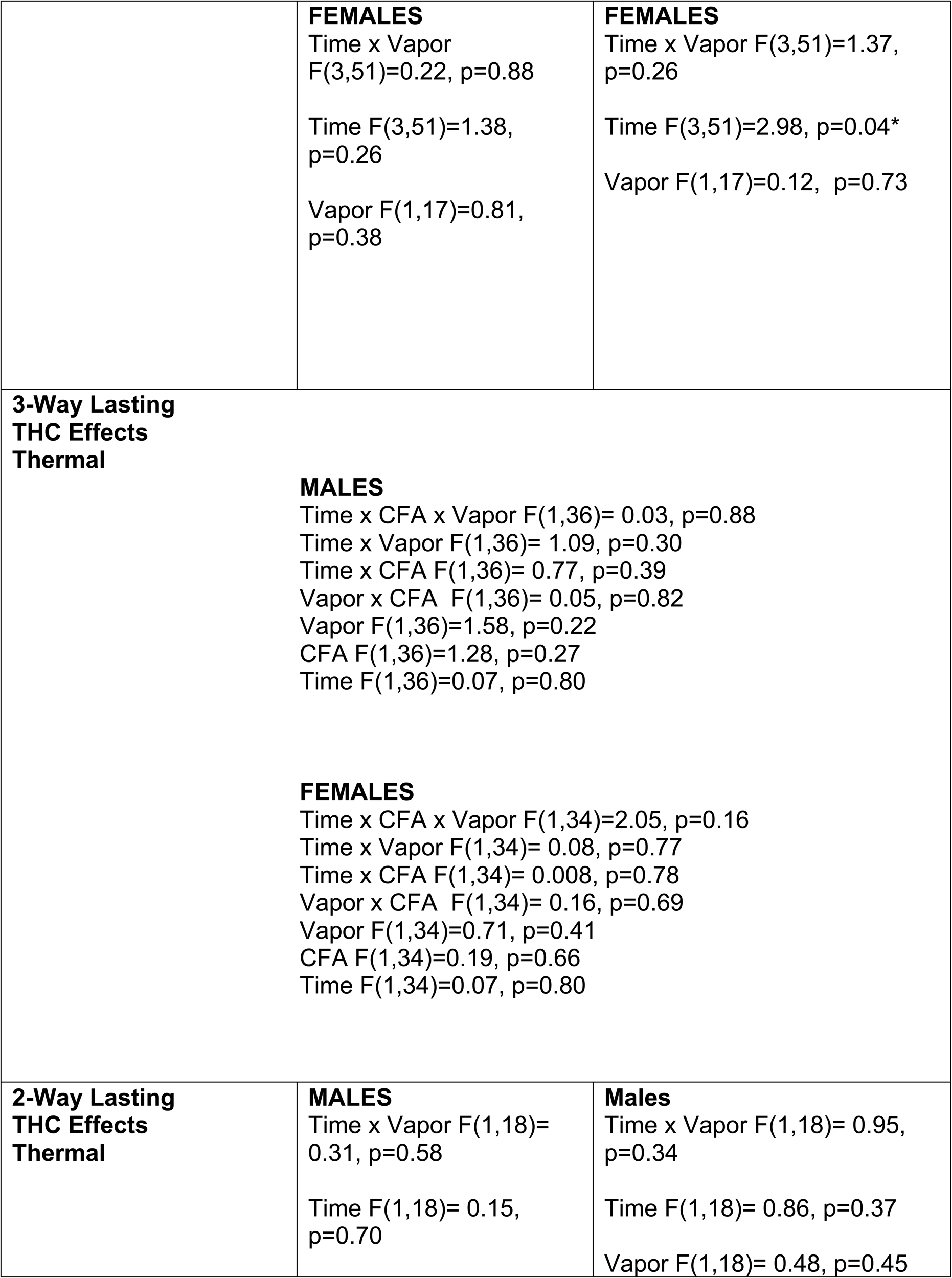
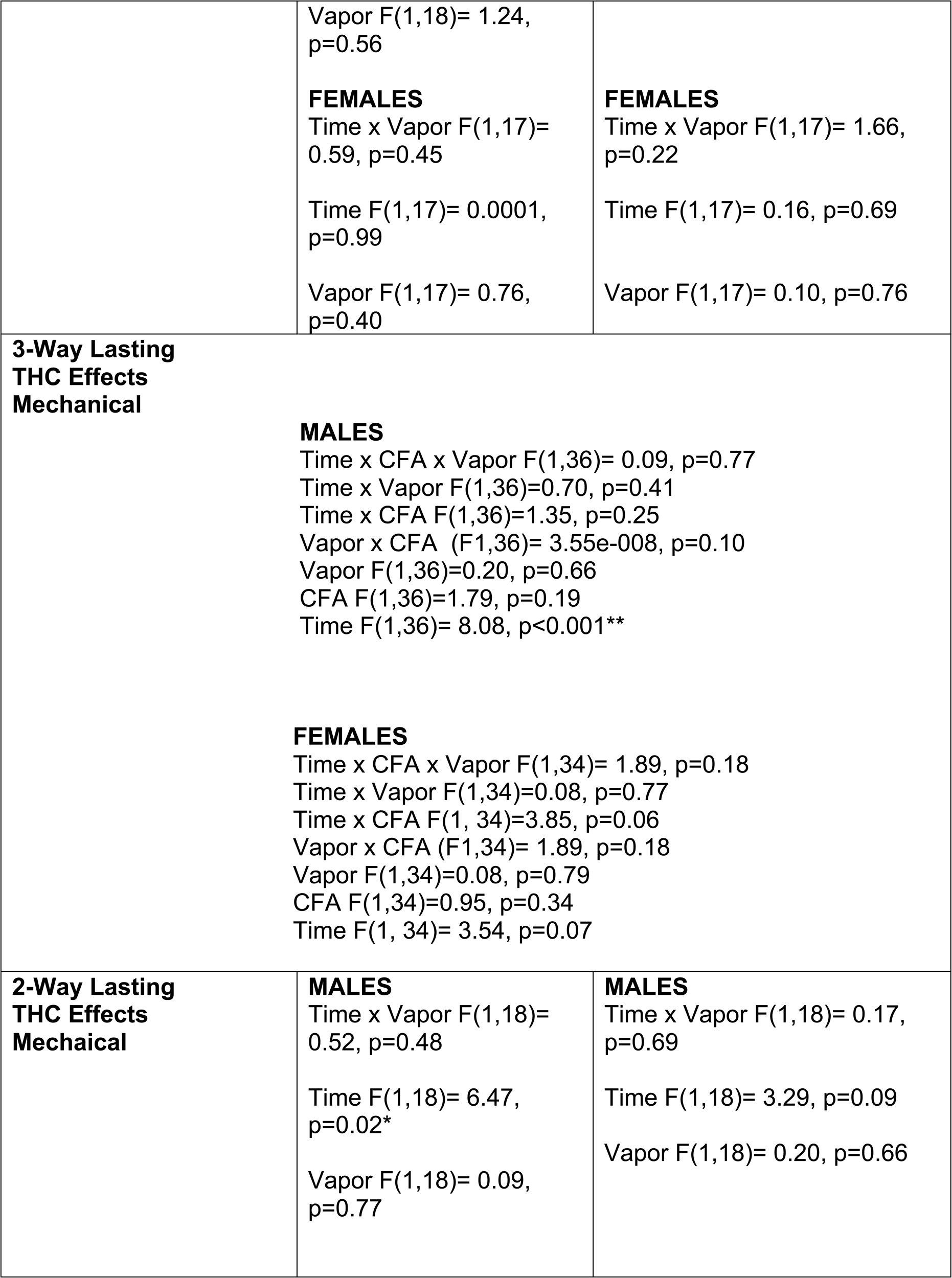
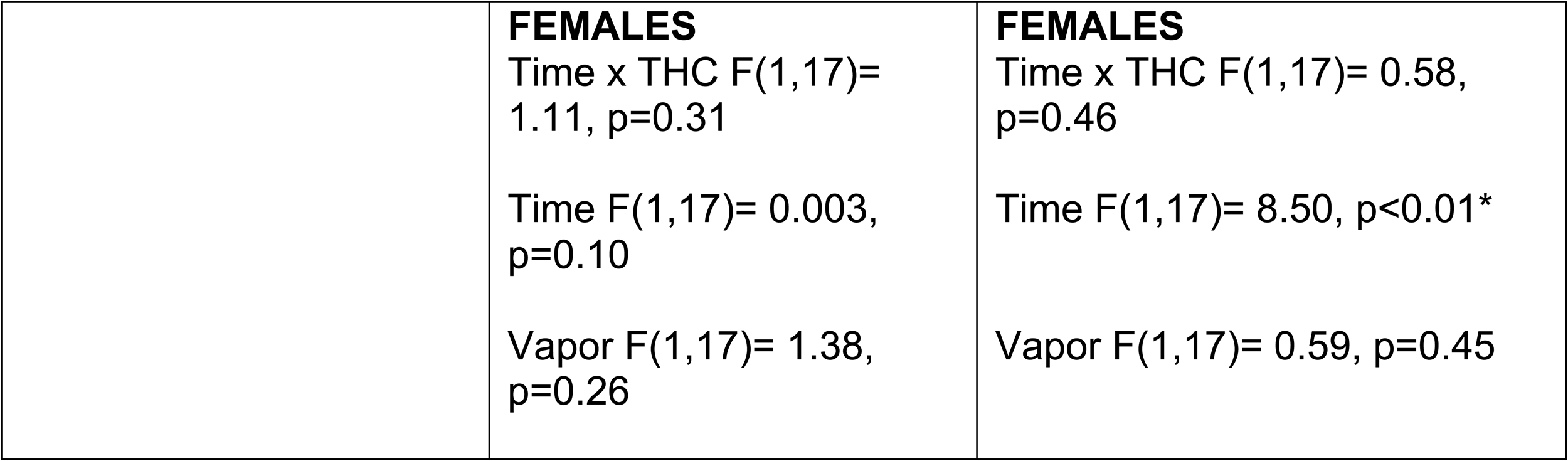
Uninjured paw thermal and mechanical data

#### Acute effects of THC vapor inhalation on mechanical sensitivity

We performed separate 3-way (CFA x vapor x time) repeated-measures (RM) ANOVAs in male and female rats for mechanical sensitivity data at baseline and acutely post-vapor exposure (**Fig. 2A**). We observed a significant main effect of CFA condition (Saline vs. CFA) in males [F(1,36)=85.64 p<0.0001] and females [F(1,34)=19.50, p<0.001], no significant effect of vapor condition (vehicle vs. THC) in males [F(1,36)=0.39, p=0.53] or females [F(1,34)=2.23, p=0.14], and a significant effect of time in females [F(3,102)=12.21, p<0.0001] but not males [F(2.57,92.42)=0.39, p=0.73]. There was a significant interaction of CFA condition x vapor condition in males [F(1,36)=4.29, p<0.05], but not in females [F(1,34)=0.45, p=0.51]. There were no other significant interactions or main effects for any other variables in males or females: time x CFA condition x vapor condition [F(3,108)=1.33, p=0.27], [F(3,102)=0.63, p=0.51]; time x vapor condition [F(3,108)=1.63, p=0.19], [F(3,102)=1.00, p=0.40]; or time x CFA condition [F(3,108)=0.43, p=0.73], [F(3,102)=0.41, p=0.75]. An analysis of the uninjured right paw revealed a significant interaction of time x vapor condition for males [F(3,108)=3.18, p=0.03], and a significant main effect of time [F(3, 102)=3.81 p=0.01) for Females. There were no other main effects or interaction effects in uninjured right paw sensitivity (**Table 1**). Tukey’s post-hoc analyses revealed that CFA-treated males exhibited higher left paw mechanical sensitivity relative to saline-treated males (p<0.05) in all comparisons.

We used separate 2-way RM ANOVAs to analyze mechanical sensitivity data collected in <u>CFA-treated male and female rats</u> over days of vaporized THC or vehicle exposure (**Fig. 2A**). THC vapor exposure increased the force required to produce a (left) hindpaw withdrawal in males [main effect of vapor; [F(1,18)=5.68, p=0.03], but not in females [F(1,17)=3.29, p=0.09]. There was a significant main effect of time in females [F(3,51)=9.90, p<0.0001] but not in males [F(3,5)=0.26, p=0.86], and there was no significant vapor x time interaction effect in Males [F (3,54)=0.21, p=0.89] or Females [F(3,51)=0.46, p=0.71].

We used separate 2-way RM ANOVAs to analyze mechanical sensitivity data collected in <u>saline-treated male and female rats</u> over days of vaporized THC or vehicle exposure (**Fig. 2A**). There were no significant effects of THC vapor treatment in males [F(1,18)=0.77, p=0.39] or females [F(1,17)=0.26, p=0.61]. There was a significant main effect of time in females [F(3,51)=3.76, p=0.02], but not in males [F (2.26,90.63)=0.47, p=0.65], and there was no vapor x time interaction effect in males [F (3,54)=1.97, p=0.13] or females [F(3,51)=1.07, p=0.37]. There was a main effect of time for the uninjured right paw in CFA females [F(3,51)=2.98, p=0.04], but no other interactions or main effects in uninjured right paw sensitivity (**Table 1**).

#### Lasting effects of THC vapor inhalation on mechanical sensitivity

We performed separate 3-way (CFA x vapor x time) repeated-measures (RM) ANOVAs in male and female rats for mechanical sensitivity data at baseline and 24 hours after the last THC or vehicle vapor exposure (**Fig. 2A**). There were no significant 3-way interaction effects in males [F(1,36)=3.29, p=0.08] or females [F(1,34)=3.30, p=0.10]. There were also not significant 2-way time x vapor interaction effects in males [F(1,36)=0.68, p=0.42] or females [F(1,34)=0.14, p=0.71], and no significant time x CFA interaction effects in males [F(1,36)=2.08, p=0.16] or females [F(1,34)=3.24, p=0.08]. There was a significant interaction of vapor x CFA condition in males [F(1,36)=7.23, p<0.05], but not females [F(1,34)=0.00013, p=0.99]. Finally, there was a significant main effect of CFA condition in males [F(1,36)=125.70, p<0.001] and females [F(1,34)=14.07, p<0.001], no significant main effect of vapor condition in males [F(1,36)=0.08, p=0.78] or females [(1,34)=1.76, p=0.19], and a significant main effect of time in females [F(1,34)=5.01, p=0.03], but not in males [F(1,36)= 0.52, p=0.48]. There was a significant main effect of time for the uninjured right paw of males [F(1,36)= 8.08, p<0.01], but no other significant main effects or interaction effects in uninjured right paw sensitivity (**Table 1**). Tukey’s post hoc analysis revealed that CFA-treated males had higher mechanical sensitivity compared to saline-treated males (p<0.0001) at both timepoints.

We used separate 2-way RM ANOVAs to analyze mechanical sensitivity data collected in <u>CFA-treated male and female rats</u> 24 hours after the last THC or vehicle vapor exposure versus baseline (**Fig. 2A**). THC vapor inhalation reduced mechanical sensitivity relative to pre-vapor baseline in CFA-treated males [F(1,18)= 7.43, p=0.01], but not in CFA-treated females [F(1,17)=0.89, p=0.36]. There was a main effect of time for CFA-treated females. CFA females exhibited higher paw withdrawal thresholds 24 hours after the final vapor exposure relative to pre-vapor baseline [F(1,17)=6.93, p=0.02], but this effect was not seen in males [F(1,18)= 0.28, p=0.60]. There was no time x vapor interaction effect in males [F(1,18)=0.53, p=0.48] or females [F(1,17)=0.06, p=0.81]. There was also a significant main effect of time seen in the uninjured right paw of CFA-treated females [F(1,17)= 8.50, p=0.01], but no other significant interactions or main effects in uninjured right paw sensitivity (**Table 1**).

We used separate 2-way RM ANOVAs to analyze mechanical sensitivity data collected in <u>saline-treated male and female rats</u> 24 hours after the last THC or vehicle vapor exposure and baseline (**Fig. 2A**). There was no main effect of THC treatment in males [F(1,18)= 2.06, p=0.17] or females [F(1,17)=0.88, p=0.36]. There was no main effect of time in males [F(1,18)= 2.19, p=0.16] or females [F(1,17)=0.12, p=0.74]. Finally, there was no time x vapor interaction effect in males [F(1,18)= 3.26, p=0.09] or females [F(1,17)=0.09, p=0.77]. There was a significant main effect of time in the uninjured right paw of saline treated males [F(1,18)=6.47, p=0.02], but no other significant interactions or main effects in uninjured right paw sensitivity (**Table 1**).

#### Acute effects of THC vapor inhalation on thermal nociception

Separate 3-way RM ANOVAs of thermal nociception data (**Fig. 2B**) for saline- and CFA-treated male and female rats following exposure to vaporized THC or vehicle exposure revealed no significant interaction of CFA condition x vapor condition in males [F(1,36)=3.91, p=0.06] or females [F(1,34)=2.64, p=0.11], a significant interaction of time x vapor condition in both males [F(3,108)=6.06, p=0.001] and females [F(3,102)=2.96, p=0.04], and a significant interaction of time x CFA condition in males [F(3,108)=8.84, p<0.0001] but not females [F(3,12.71)=2.25, p=0.09]. There was also a significant main effect of CFA condition for males [F(1,36)=51.28, p<0.0001] and females [F(1,34)=11.69, p=0.002], a significant main effect of vapor condition for males [F(3,108)=6.06, p<0.001] and females [F(1,34)=5.46, p=0.03], and a significant main effect of time in males [F(2.76,99.39)=8.45, p<0.0001], but not in females [F(3,7.54)=1.34, p=0.27]. There was no significant CFA x vapor x time 3-way interaction in males [F(3,108)=1.38, p=0.25] or females [F(3,11.98)=2.12, p=0.10]. There was a significant main effect of vapor on thermal nociception in the uninjured right paw of males [F(1,36)=4.88, p=0.03], and significant main effects of time for both males [F(2.58,92.71)=3.48, p=0.02] and females [F(2.89, 98.12)=3.453, p=0.02]. There were no other main effects or interaction effects of CFA condition or vapor condition on thermal nociception in the uninjured right paw (**Table 1**). Tukey’s post-hoc test revealed that CFA-treated male rats exhibited thermal hyperalgesia in the injured paw at baseline relative to saline-treated male rats (p<0.05). THC vapor rescued thermal hyperalgesia in CFA-treated male rats on day 2 (p=0.02) and day 5 (p=0.03) relative to vehicle-treated CFA male rats. THC also rescued thermal hyperalgesia in in CFA-treated male rats on day 2 (p=0.02) and day 5 (p=.04) relative to their own baseline. Tukey’s post-hoc test revealed that CFA induced thermal hyperalgesia in female rats relative to saline-treated females (p<0.05), and that THC treatment rescued thermal hyperalgesia in CFA-treated female rats on day 2 (p=0.05).

Separate 2-way RM ANOVAs of thermal nociception data in <u>CFA-treated male and female rats</u> revealed that THC vapor exposure rescued CFA-induced thermal hyperalgesia in both males and females (**Fig. 2B**). Statistical analysis confirmed a significant main effect of time in males [F(2.45,44.02)=19.69 p<0.0001] but not females [F(3,51)=2.35, p=0.08], a significant main effect of THC treatment in males [F(1,18)=28.75, p<0.0001] and females [F(1,17)=5.96, p=0.03], and a significant vapor x time interaction effect in Males [F(3,54)=7.22, p<0.01] and Females [F(3,51)=3.93, p=0.01]. Post-hoc tests confirmed that THC (100 mg) vapor exposure compared with PG vehicle increased paw withdrawal latency in male CFA-treated rats on vapor days 2, 5 and 8 (p<0.05) and female CFA-treated rats on vapor days 2 (p<0.01) and 8 (p<0.05). There was no main effect of vapor condition on thermal nociception in the uninjured right hindpaw (**Table 1**).

Separate 2-way RM ANOVAs of thermal nociception data in <u>saline-treated male and female rats</u> revealed that THC vapor exposure did not significantly alter thermal nociception relative to vehicle-treated male or female rats (**Fig. 2B**). Statistical analysis showed no significant main effects of time in males [F(2.26,90.63)=0.47, p=0.65] or females [F(3,51)=1.11, p=0.35], no significant main effect of THC treatment in males [F (THC F(1,18)=3.96, p=0.06] or females [F(1,17)=0.55, p=0.47], and no vapor x time interaction effects in males [F(3,54)=1.97, p=0.13] or females [F(3,51)=0.97, p=0.42]. There was no main effect of vapor condition on nociception in the uninjured right hindpaw (**Table 1**).

#### Lasting effects of THC vapor inhalation on thermal nociception

Separate 3-Way RM ANOVAs of thermal nociception data in male and female rats at 24 hours after the last exposure to THC or vehicle vapor and baseline (**Fig. 2B**) showed a significant interaction of time x CFA condition for males [F(1,36)=13.55 p<0.001], but not females [F(1,34)=3.23, p=0.08], and a significant interaction for time x vapor condition in females [F(1,34)=7.64, p<0.01], but not males [F(1,36) p=0.17]. There was no significant interaction for CFA condition x vapor condition in males [F(1,36)=2.21, p=0.15] or females [(1,34)=1.41, p=0.24]. There was a significant main effect of CFA condition in males [F(1,36)=91.99, p<0.0001] and females [F(1,34)=14.18, p<0.001], a significant main effect of vapor condition in males [F(1,36)=4.0, p=0.05], but not females [F(1,34)=3.52, p=0.06], and a significant main effect of time in males [F(1,36)=26.39, p<0.001] and females [F(1,34)=9.92, p<0.01). There were no significant 3-way interactions for time x CFA condition x vapor condition in male rats [F(1,36)=1.94, p=0.17] or female rats [F(1,34)=0.54, p=0.47]. There were no main effects or interaction effects of CFA condition and vapor condition on nociception in the uninjured right hindpaw (**Table 1**). Tukey’s post-hoc test revealed that CFA induced thermal hyperalgesia in the injured paw of male rats relative to saline-treated males (p<0.0001), and that THC rescued thermal hyperalgesia in CFA-treated male rats 24 hours after the last THC vapor session relative to baseline (p=0.002) and also relative to vehicle-treated CFA males (p<0.05). Tukey’s post-hoc test revealed that CFA induced thermal hyperalgesia in the injured paw of CFA-treated females relative to saline-treated females (p<0.05), and that THC rescued thermal hyperalgesia in CFA-treated females 24 hours after the last THC vapor session relative to baseline (p<0.01) and also relative to vehicle-treated CFA females (p<0.05).

Separate 2-way RM ANOVAs of thermal nociception data collected in <u>CFA-treated male and female rats</u> 24 hours after the last vapor exposure versus baseline (**Fig. 2B**) showed a significant main effect of THC treatment in males [THC F(1,18)=6.78, p=0.02] but not females [F(1,17)=3.94 p=0.06], a significant main effect of time in males [F(1,18)=28.78, p<0.001] and females [F(1,17)=9.38, p<0.01], and a significant THC x Time interaction effect in females [F(1,17)=6.59, p=0.02] but not males [F(1,18)= 2.88, p=0.11]. Tukey’s post-hoc analysis revealed that THC rescued hyperalgesia in CFA-treated females (p<0.01) 24 hours after the last vapor exposure relative to vehicle-exposed CFA females. There was no main effect of vapor condition on nociception in the uninjured right hindpaw (**Table 1**).

Separate 2-way RM ANOVAs of thermal nociception data collected in <u>saline-treated male and female rats</u> 24 hours after the last vapor exposure versus baseline (**Fig. 2B**) showed no significant effects of THC treatment in males [F(1,18)= 0.20, p=0.73] or females [F(1,17)=0.30, p=0.60], no significant effect of time in males [F(1,18)= 1.63, p=0.22] or females [F(1,17)=1.32, p=0.27] and no vapor x time interaction effect in males [F(1,18)= 9.703e-007, p=0.99] or females [F(1,17)=1.37, p=0.26]. There were no significant main effects or interaction effects on nociception in the uninjured right hindpaw (**Table 1**).

### Experiment 2

#### Plasma THC levels following THC vapor exposure days 1 and 10

We used a 2-way RM ANOVA to analyze THC and metabolite levels in plasma samples collected from CFA- and saline-treated <u>males</u> on days 1 and 10 following exposure to 100 mg THC vapor (**Fig. 3**). There were no significant main effects of CFA condition [F(1,12)=0.396, p=0.54] or time [F(1,12)=3.618, p=0.08] and no time x CFA interaction effects on plasma THC levels [F(1,12)=0.639, p=0.44]. There were no significant main effects of CFA condition [F(1,12)=0.423, p=0.53] or time [F(1,12)= 0.101, p=0.76] and no time x CFA interaction effects on 11-OH-THC levels in males [F(1,12)=0.053, p = 0.82]. There was no significant main effect of CFA condition [F(1,12)=0.168, p=0.69] and no time x CFA interaction effect on THC-COOH levels in males [F(1,12)=0.955, p=0.35], but there was a significant main effect of time on THC-COOH levels in males [F(1,12)=8.374, p=0.01].

**Figure 3:**
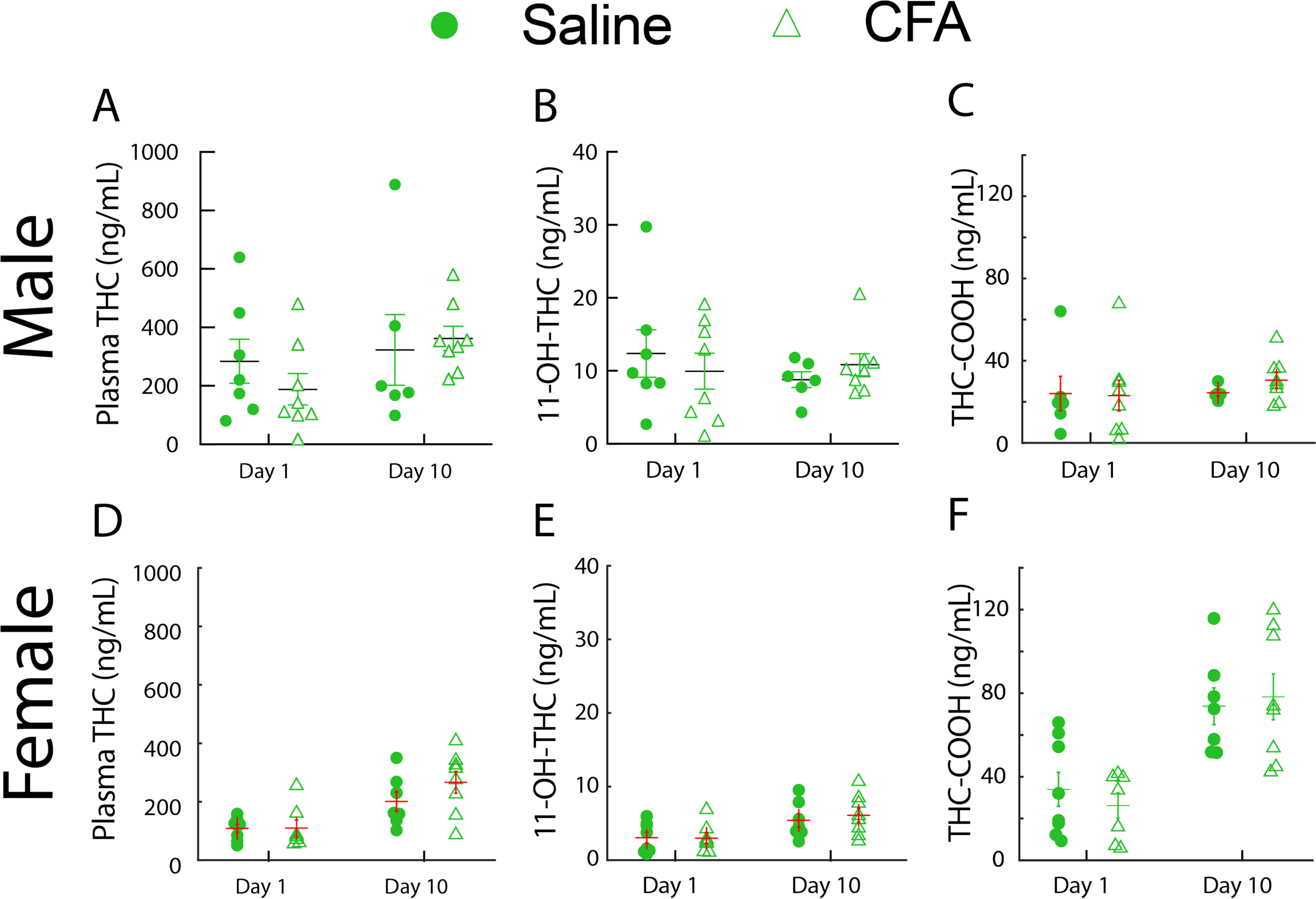
Plasma THC and THC metabolites Analysis following vapor exposure days 1 and 10. (**A**) A mixed effects analysis of plasma samples collected from CFA and Saline treated rats on days 1 and 10 following exposure to 100 mg THC vapor (Fig. 3) showed no significant effects of CFA condition [F(1,25)=0.1528 p=0.6991] or Time [F(1,25)=2.164 p=0.1538] and no interaction of factors [F(1,25)=0.8779 p=0.3577] in THC plasma levels. Analysis of THC metabolites showed no significant effects of Pain condition [F(1,13)=0.008842 p=0.9265] or Time [F(1,12)= 0.3816 p=0.5483] and no interaction of factors [F(1,12)=1.085] on 11-OH-THC levels and no significant effects of CFA condition [F(1,13)=0.02756 p=0.8707] or Time [F(1,12)=1.451 p=0.2516] or interaction of factors [F(1,12)=0.1838 p=0.6757] on THC-COOH levels.

We used a separate 2-way RM ANOVA to analyze THC and metabolite levels in plasma samples collected from CFA- and saline-treated <u>females</u> on days 1 and 10 following exposure to 100 mg THC vapor (**Fig. 3**). There was a significant main effect of time [F(1,12)=35.22, p<0.0001], but no main effect of CFA condition [F(1,12)=0.499, p<0.50] and no time x CFA interaction effect on plasma THC levels [F(1,12)=2.207, p<0.16]. There was a significant main effect of time [F(1,12)=8.38, p=0.01], but no main effect of CFA condition [F(1,12)=0.430, p=0.79] and no time x CFA interaction effects on 11-OH-THC metabolite levels in females [F(1,12)=0.312, p=0.59]. Finally, there was a significant main effect of time [F(1,12)=28.067, p<0.001], but no main effect of CFA condition [F(1,12)=0.052, p=0.82] and no time x CFA interaction effect on THC-COOH metabolite levels in females [F(1,14)=0.818, p=0.38].

We performed a 2-way (sex x time) RM ANOVA to compare plasma levels of THC and its metabolites in male and female rats; CFA and saline data were pooled because there were no main effects of CFA on any measured plasma-drug or plasma-metabolite levels. We observed a significant main effect of sex [F(1,26)=5.399, p=0.03] and a significant main effect of time [F(1,26)=14.059, p<0.001], but no time x sex interaction effect [F(1,26) = 0.03, p. = 0.86] on plasma-THC levels. There was a significant main effect of sex [F(1,26)=11.697, p<0.01] on 11-OH-THC levels in plasma, but no main effect of time [F(1,26)=3.277, p=0.08] and no time x sex interaction effect [F(1,12)=1.376, p=0.25]. Finally, we observed a main effect of sex [F(1,26)=11.086, p<0.01], a main effect of time [F(1,26)=36.141, p<0.0001], and a significant time x sex interaction effect on plasma THC-COOH levels [F(1,26)=13.593, p=0.001].

### Experiment 3

#### CFA and THC effects on intrinsic properties in all vlPAG neurons

We tested the effects of CFA and chronic THC vapor inhalation on neural and synaptic properties in vlPAG neurons by performing whole cell recordings in male and female rats. Voltage and current recordings were performed to characterize measures of intrinsic excitability: resting membrane potential [RMP], input resistance (R_input_) and voltage SAG. A 2-way ANOVA for RMP data revealed no effect of CFA [F(1,150) = 0.002, p = 0.97] or vapor [F(1,150) = 0.14, p = 0.71], and no interaction effect [F(1,150) = 0.03, p = 0.86] (**Fig. 4A**) in males. A 2-way ANOVA for R_input_ data showed no main effect of CFA [F(1,166) = 0.2, p = 0.65] or vapor [F(1,166) = 2.1, p = 0.15], and no interaction effect [F(1,166) = 0.96, p = 0.33] (**Fig. 4B**) in males. Figure 4C shows the characteristic rebound voltage excursion seen in some neurons following hyperpolarization caused by negative current steps, known as voltage SAG. A 2-way ANOVA for SAG data showed no main effect of CFA [F(1,173) = 0.3, p = 0.60] or vapor [F(1,173) = 2.3, p = 0.13], and no interaction effect [F(1,173) = 3.73, p = 0.055] (**Fig. 4D**) in males. Figure 4E shows membrane voltage responses to current step inputs. Figure 4F shows the FI curve constructed from the number of action potentials during each current step. 2-way ANOVA analysis of the Firing rate @ 150 pA revealed a significant main effect of vapor [F(1,174) = 4.19, p = 0.042], with lower firing rates in the THC groups and an effect size of 0.024. There was no effect of CFA [F(1,174) = 0.003, p = 0.96] or CFA x vapor interaction effect [F(1,174) = 0.54, p = 0.46] (**Fig. 4G**) in males, Tukey’s post-hoc pairwise comparisons revealed no significant differences between individual groups. A 2-way ANOVA for gain of firing rate curves, calculated between 50 and 150 pA inputs, showed no main effect of CFA [F(1,174) = 0.01, p = 0.92] or vapor [F(1,174) = 3.56, p = 0.061], and no interaction effect [F(1,174) = 0.59, p = 0.44] (**Fig. 4H**) in male rats. Tukey’s post-hoc pairwise comparisons revealed no significant differences between individual groups.

**Figure 4:**
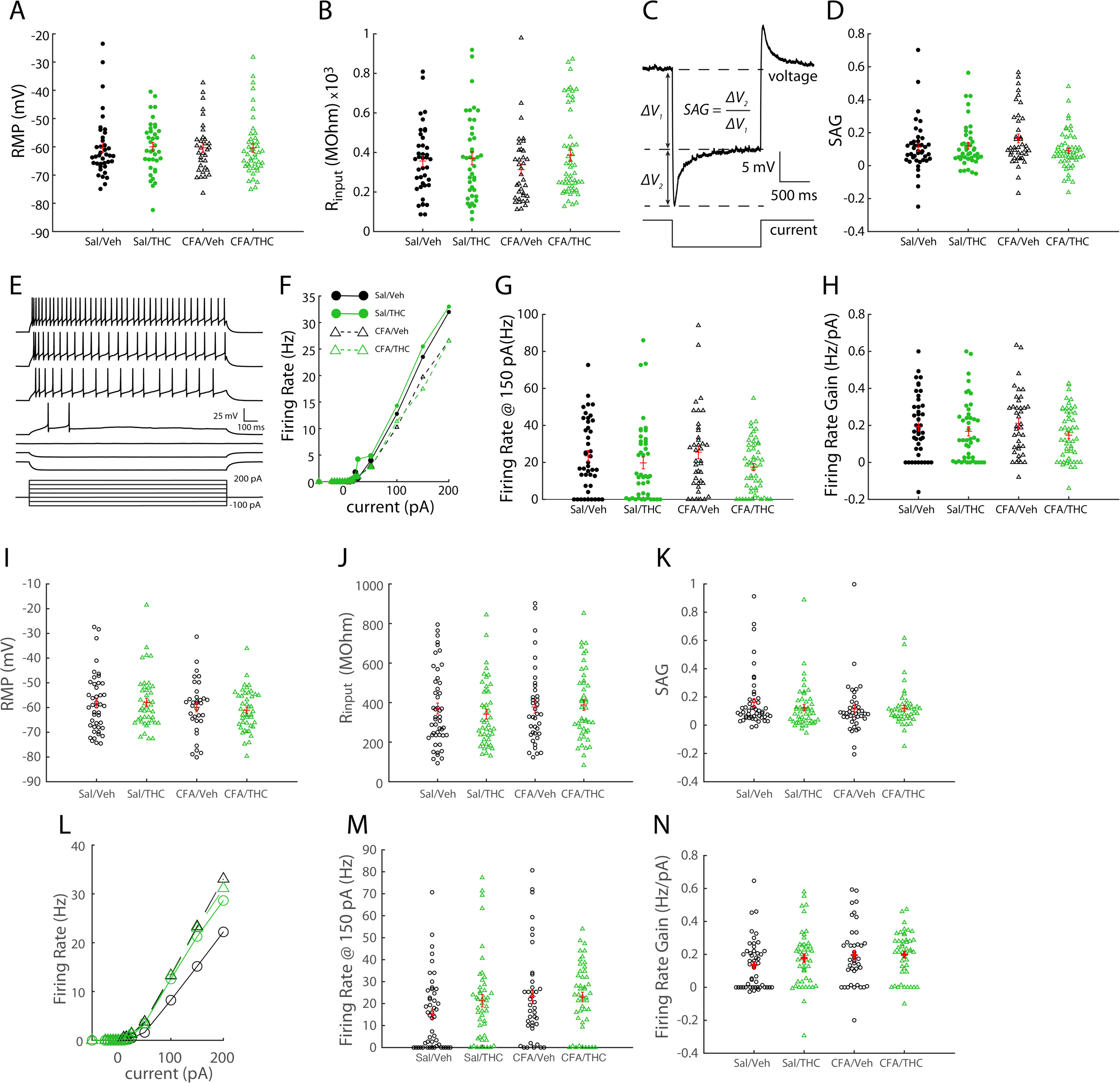
Effects of CFA and vapor on intrinsic properties of vlPAG neurons. 2-Way ANOVAs in male rats show: (**A**) no effect of CFA [F(1,150) = 0.002, p = 0.9686] or vapor [F(1,150) = 0.14, p = 0.7062] on RMP, (**B**) no effect of CFA [F(1,166) = 0.2, p = 0.6522] or vapor [F(1,166) = 2.1, p = 0.1489] on R_input_, and (**C,D**) no effect of CFA [F(1,166) = 0.2, p = 0.6522] or vapor [F(1,166) = 0.96, p = 0.3277] on SAG. (**E**) An example set of voltage responses to injected current steps of different magnitudes. (**F**) Mean firing Rate response curves of vlPAG neurons to injected current. (**G**) There was a main effect of vapor on firing rate in vlPAG neurons [F(1,174) = 4.19, p = 0.0421]. (**H**) There was no main effect of CFA [F(1,174) = 0.01, p = 0.9241] or THC vapor [F(1,174) = 3.56, p = 0.061] on firing rate gain in vlPAG neurons. 2-Way ANOVAs in female rats show: (**I**) no effect of CFA [F(1,161) = 0.203, p = 0.1562] or vapor [F(1,161) = 0.05, p = 0.8223] on RMP, (**J**) no effect of CFA [F(1,171) = 0.97, p = 0.3254] or vapor [F(1,171) = 0.11, p = 0.7406] on R_input_, and (**K**) no effect of Pain [F(1,176) = 0.99, p = 0.928] or vapor [F(1, 176) = 0.37, p = 0.5417] on SAG. (**L,M**) There was no main effect of CFA [F(1,179) = 3.44, p = 0.0651] or vapor [F(1,179) = 1.19, p = 0.277] on firing rates in vlPAG neurons [F(1,174) = 4.19, p = 0.0421]. (**N**) There was no main effect of CFA [F(1,179) = 2.9, p = 0.0902] or THC vapor [F(1, 179) = 0.89, p = 0.3461] on firing rate gain in vlPAG neurons.

In females, a 2-way ANOVA for RMP data showed no effect of CFA [F(1,161) = 2.03, p = 0.16] or vapor [F(1,161) = 0.05, p = 0.82], and no interaction effect [F(1,161) = 0.26, p = 0.61] (**Fig. 4I**). A 2-way ANOVA for R_input_ data showed no main effect of CFA [F(1,171) = 0.97, p = 0. 33] or vapor [F(1, 171) = 0.11, p = 0.74], and no interaction effect [F(1, 171) = 0.49, p = 0.49] (**Fig. 4J**) in females. A 2-way ANOVA for SAG data showed no main effect of CFA [F(1,176) = 0.99, p = 0.32] or vapor [F(1, 176) = 0.37, p = 0.5417], and no interaction effect [F(1, 176) = 0.77, p = 0.38] (**Fig. 4K**) in females. A 2-way ANOVA for firing rate @ 150 pA revealed a near-significant main effect of CFA [F(1,179) = 3.44, p = 0.065] (**Fig. 4M**), in females. There was no effect of THC [F(1, 179) = 1.19, p = 0.28] or CFA x vapor interaction effect [F(1, 179) = 1.46, p = 0.23]. A 2-way ANOVA for gain of the firing rate curves, calculated between 50 and 150 pA inputs, showed no main effect of CFA [F(1, 179) = 2.9, p = 0.090] or vapor [F(1, 179) = 0.89, p = 0.35], and no interaction effect [F(1, 179) = 0.67, p = 0.41] (**Fig. 4N**) in females.

#### CFA and THC effects on synaptic properties in all vlPAG neurons

Voltage clamp recordings were performed to characterize synaptic properties of vlPAG neurons: with blockers of glutamate receptors (APV [50 μM] and CNQX [10 μM]) we recorded postsynaptic currents while clamping the voltage at -50 mV (**Fig. 5A**) and detected unitary spontaneous inhibitory current events (**Fig. 5B**) in males. The amplitude and frequency of these events was calculated to assess net inhibitory inputs to the recorded vlPAG neurons. 2-way ANOVA for sIPSC amplitude data revealed a significant main effect of vapor [F(1,150) = 31.91, p = 0.0003], with lower sIPSC amplitudes in the THC groups and an effect size of 0.085. There was no main effect of CFA [F(1,150) = 0.1, p = 0.75], and no CFA x THC interaction effect [F(1,150) = 0.5, p = 0.48] (**Fig. 5C**) in males. Tukey’s post-hoc pairwise comparisons revealed significant differences between Sal/Veh and Sal/THC groups (p < 0.01) and Sal/THC and CFA/Veh groups (p <0.05), with the means of the Sal/Veh and CFA/Veh groups being significantly larger than the Sal/THC group mean. A 2-way ANOVA for sIPSC frequency data revealed a main effect of vapor [F(1,150) = 8.23, p = 0.0047], with lower sIPSC frequencies in the THC groups and an effect size of 0.052. There was no main effect of CFA [F(1,150) = 0.23, p = 0.63] and no CFA x THC interaction effect [F(1,150) = 8e^-5^ p = 0.99] (**Fig. 5D**) in males. Tukey’s post-hoc pairwise comparisons revealed no significant differences between individual groups. A 2-way ANOVA for sIPSC amplitude data in female rats revealed a significant main effect of CFA [F(1,172) = 5.25, p = 0.041] with an effect size of 0.025. There was no main effect of THC [F(1, 172) = 2.4, p = 0.12]. There was also a significant CFA x THC interaction effect [F(1, 172) = 7.1, p = 0.0084] (**Fig. 5E**) in females, with an effect size of 0.041. Tukey’s post-hoc pairwise comparisons revealed significant differences between Sal/Veh and CFA/Veh groups (p < 0.01) and CFA/Veh and CFA/THC groups (p <0.05), with the means of the CFA/Veh group being significantly larger than the Sal/Veh and CFA/THC group means. A 2-way ANOVA for sIPSC frequency data showed no main effect of CFA [F(1,172) = 0.56, p = 0.4543], no main effect of THC [F(1, 172) = 0.06, p = 0.81] and no CFA x THC interaction effect [F(1, 172) = 2.46, p = 0.12] (**Fig. 5F**) in females. Post-hoc pairwise comparisons revealed no significant differences between individual groups.

**Figure 5:**
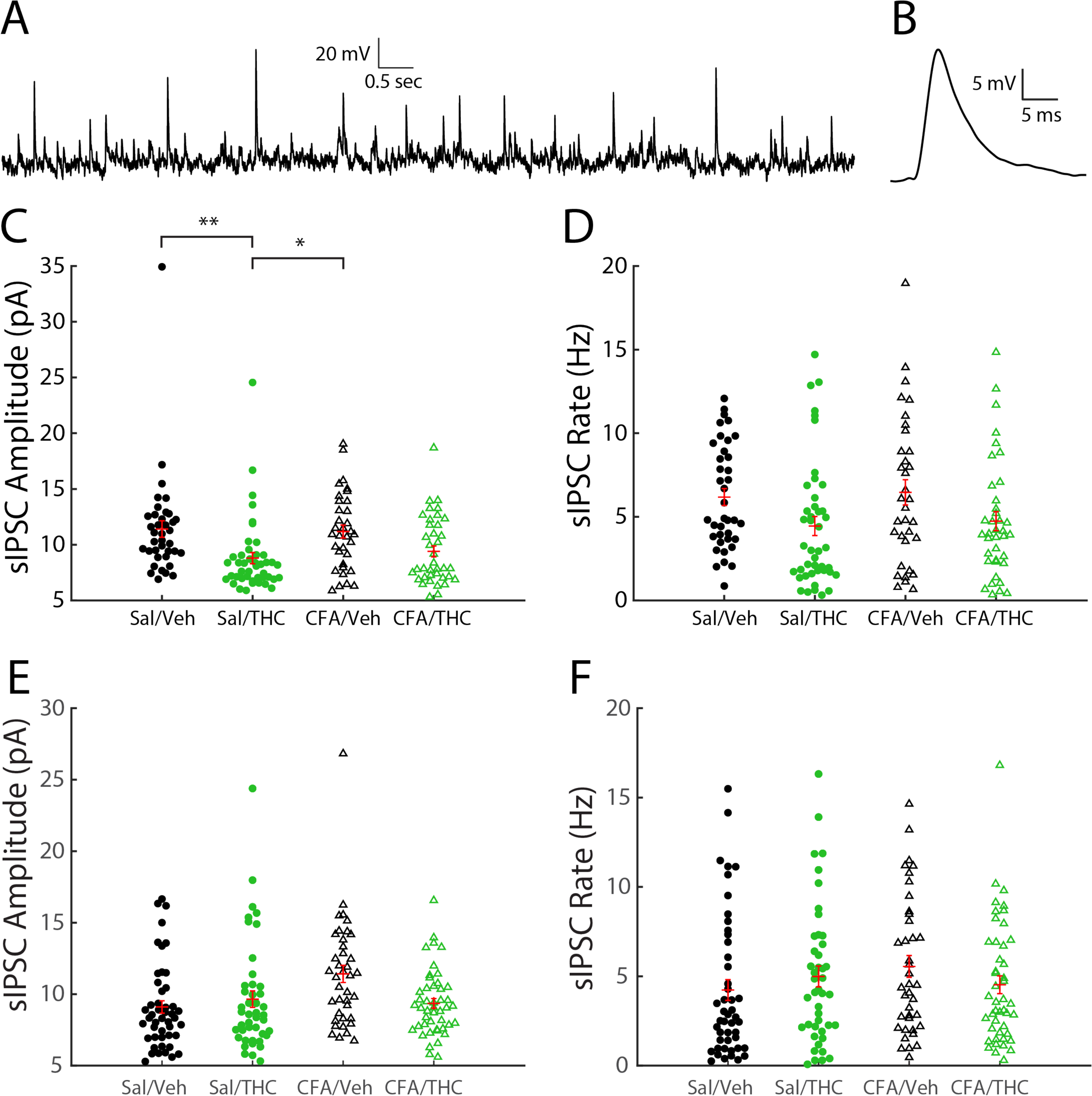
Effects of CFA and vapor of synaptic properties in vlPAG neurons. (**A**) An example current traces from a vlPAG neuron with membrane voltage clamped at -50 mV. (**B**) An example average inhibitory postsynaptic current event trace. (**C**) 2-Way ANOVA revealed a main effect of vapor on sIPSC amplitude [F(1,150) = 31.91, p = 0.0003] in vlPAG neurons from male rats. (**D**) There was also a main effect of vapor on sIPSC rate [F(1,150) = 8.23, p = 0.0047]. (**E**) 2-Way ANOVA revealed a main effect of CFA on sIPSC amplitude [F(1,172) = 5.25, p = 0.0406] in vlPAG neurons from female rats. (**F**) There was also a CFA x vapor effect on sIPSC amplitude [F(1,172) = 7.1, p = 0.0084]. *p < 0.05, **p < 0.01. There was no main effect of CFA [F(1,172) = 0.56, p = 0.4543] or vapor [F(1,172) = 0.06, p = 0.8143] on sIPSC rates in vlPAG neurons from female rats.

#### CFA and THC effects on action potential firing patterns in vlPAG neurons

We investigated action potential firing patterns to examine if there were firing phenotype-specific effects of THC vapor on the parameters investigated, particularly firing rate response curves and synaptic properties. We found that recordings were easily divided based on the delay of first AP from current step onset. One population exhibited a short latency delay from current step onset (labeled ‘Onset’ neurons) while another displayed a distinctive longer latency delay from current step onset (labeled ‘Delayed’ neurons) (**Fig. 6A**). There were no significant differences in the proportion of these types between any of the experimental groups (**Fig. 6B**; Sal/Veh: n_onset_ = 27, n_delayed_ = 16; Sal/THC: n_onset_ = 27, n_delayed_ = 17; CFA/Veh: n_onset_ = 29, n_delayed_ = 11; CFA/THC: n_onset_ = 38, n_delayed_ = 15).

**Figure 6:**
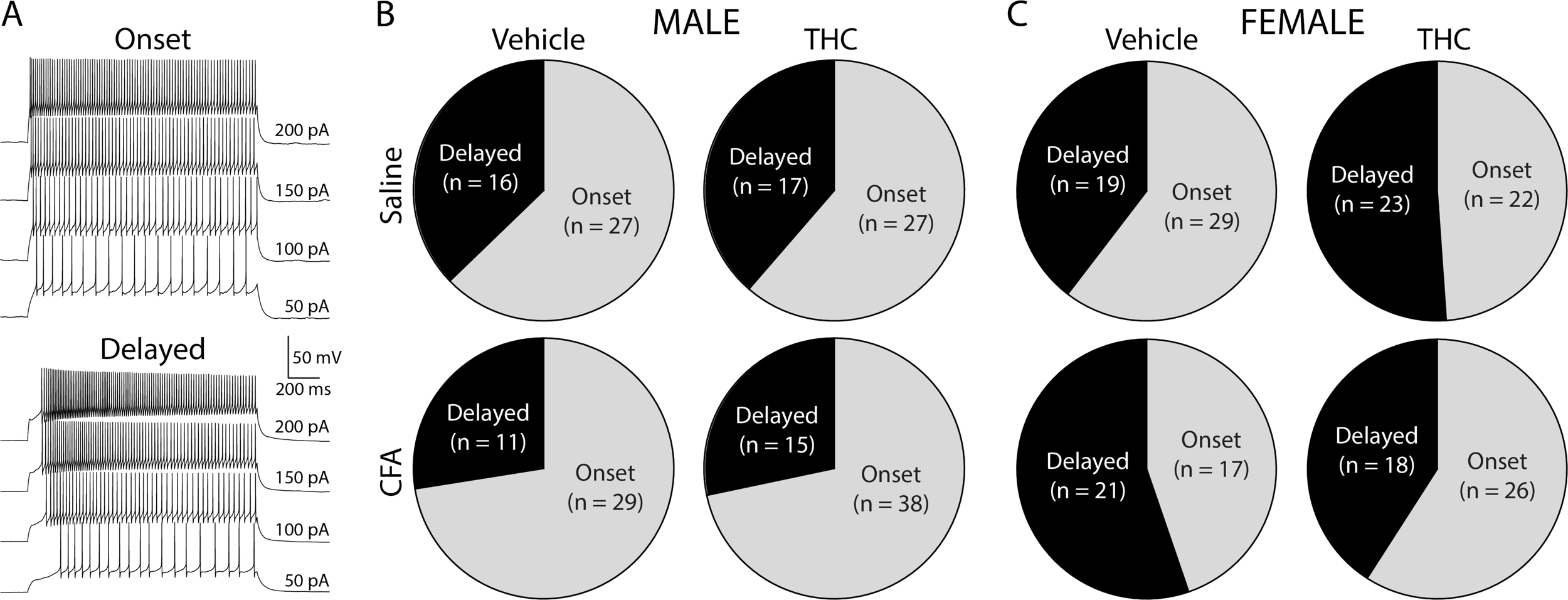
Action potential firing phenotype classification of vlPAG neurons. (**A**) Neurons in vlPAG displayed one of two different firing types based on the latency to first spike after the onset of injected depolarizing current. “Onset” neurons responded with action potentials at short latencies, while “Delayed” neurons responded with conspicuously longer latencies. (**B**) The proportion of Onset neurons (∼2/3) and Delayed neurons (∼1/3) from male rats was not significantly different for any experimental group. (**C**) The proportion of Onset neurons (∼55%) and Delayed neurons (∼45%) from female rats was not significantly different for any experimental group.

### CFA and THC effects on intrinsic properties in vlPAG Onset and Delayed neurons

In males, a 2-way ANOVA for RMP data collected from Onset neurons revealed no main effect of CFA [F(1,96) = 0.003, p = 0.96] or vapor [F(1,96) = 0.63, p = 0.43], and no CFA x THC interaction effect [F(1,96) = 1.2, p = 0.28] (**Fig. 7A**). A 2-way ANOVA for RMP data collected from Delayed neurons revealed no effect of CFA [F(1,50) = 0.06, p = 0.80] or vapor [F(1,50) = 2.03, p = 0.16], and no CFA x THC interaction effect [F(1,50) = 3.15, p =0.082] (**Fig. 7B**) in males.

**Figure 7:**
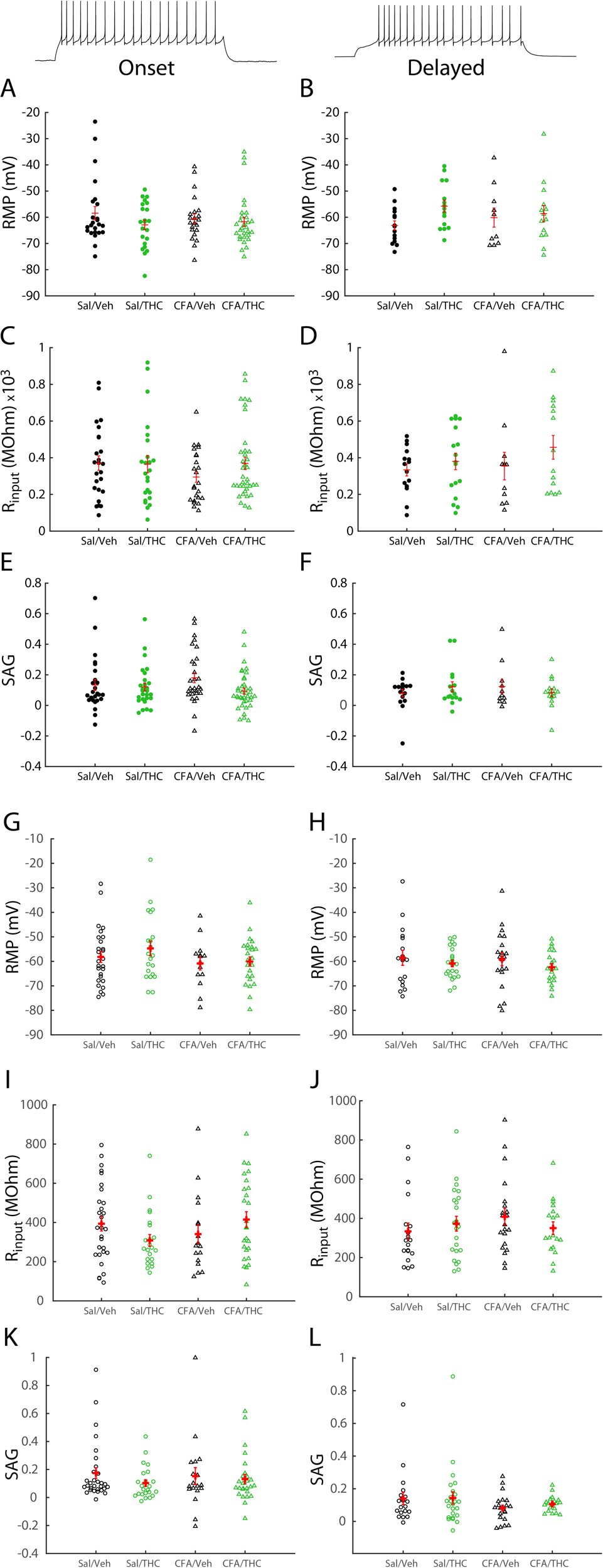
Firing phenotype-specific effects of Pain and Vapor on intrinsic properties of vlPAG neuron. (**A**) 2-Way ANOVA shows no main effect of CFA [F(1,96) = 0.003, p = 0.958] or vapor [F(1,96) = 0.63, p = 0.4286] on RMP in Onset vlPAG neurons from male rats. (**B**) There is no main effect of CFA [F(1,50) = 0.06, p = 0.8011] or vapor [F(1,50) = 2.03, p = 0.1608] on RMP in Delayed neurons. (**C**) There is no main effect of CFA [F(1,106) = 0.18, p = 0.676] or vapor [F(1,106) = 0.22, p = 0.6433] on R_input_ in Onset neurons. (**D**) There is no main effect of VFA [F(1,53) = 0.06, p = 0.8098] or vapor [F(1,53) = 3.23, p = 0.0779] on R_input_ in Delayed neurons. (**E**) There is no main effect of CFA [F(1,112) = 0.26, p = 0.6122] or vapor [F(1,112) = 2.47, p = 0.1186] on SAG in Onset neurons. (**F**) There is no main effect of CFA [F(1,54) = 0.85, p = 0.3594] or vapor [F(1,54) = 0.1, p = 0.751] on SAG in Delayed neurons. (**G**) 2-Way ANOVA shows no main effect of CFA [F(1,83) = 2.53, p = 0.1158] or vapor [F(1,83) = 0.71, p = 0.4015] on RMP in Onset vlPAG neurons from female rats. (**H**) There is no main effect of CFA [F(1,74) = 0.24, p = 0.6282] or vapor [F(1,74) = 1.59, p = 0.2106] on RMP in Delayed neurons. (**I**) There is no main effect of CFA [F(1,90) = 0.45, p = 0.5033] or vapor [F(1,90) = 0.02, p = 0.886] on R_input_ in Onset neurons. There is a CFA x vapor interaction effect [F(1,90) = 4.01, p = 0.0482] on R_input_. (**J**) There is no main effect of CFA [F(1,77) = 0.43, p = 0.5137] or vapor [F(1,77) = 0.06, p = 0.8074] on R_input_ in Delayed neurons. (**K**) There is no main effect of CFA [F(1,93) = 0.01, p = 0.928] or vapor [F(1,93) = 1.35, p = 0.2483] on SAG in Onset neurons. (**L**) There is no main effect of CFA [F(1,79) = 2.64, p = 0.1084] or vapor [F(1,79) = 0.22, p = 0.7155] on SAG in Delayed neurons.

In males, a 2-way ANOVA for R_input_ data collected from Onset neurons revealed no main effect of CFA [F(1,106) = 0.18, p = 0.68] or vapor [F(1,106) = 0.22, p = 0.64], and no CFA x THC interaction effect [F(1,106) = 0.37, p = 0.54] (**Fig. 7C**). A 2-way ANOVA for R_input_ data collected from Delayed neurons revealed no main effect of CFA [F(1,53) = 0.06, p = 0.81] or vapor [F(1,53) = 3.23, p = 0.078], and no CFA x THC interaction effect [F(1,53) = 0.67, p = 0.42] (**Fig. 7D**) in males.

In males, a 2-way ANOVA analysis of SAG data collected from Onset neurons revealed no main effect of CFA [F(1,112) = 0.26, p = 0.61] or vapor [F(1,112) = 2.47, p = 0.12], and no CFA x THC interaction effect [F(1,112) = 1.29, p = 0.26] (**Fig. 7E**). A 2-way ANOVA analysis of SAG data collected from Delayed neurons revealed no main effect of CFA [F(1,54) = 0.85, p = 0.36] or vapor [F(1,54) = 0.1, p = 0.75], and no interaction effect [F(1,54) = 3.08, p = 0.085] (**Fig. 7F**) in males.

In females, a 2-way ANOVA analysis of RMP data collected from Onset neurons revealed no main effect of CFA [F(1, 83) = 2.53, p = 0.12] or vapor [F(1, 83) = 0.71, p = 0.40], and no CFA x THC interaction effect [F(1, 83) = 0.28, p = 0.60] (**Fig. 7G**). A 2-way ANOVA analysis of RMP data collected from Delayed neurons revealed no effect of CFA [F(1,74) = 0.24, p = 0.63] or vapor [F(1, 74) = 1.59, p = 0.21], and no CFA x THC interaction effect [F(1, 74) = 0.05, p =0.82] (**Fig. 7H**) in females.

In females, a 2-way ANOVA for R_input_ data collected from Onset neurons revealed no main effect of CFA [F(1,90) = 0.45, p = 0.50] or vapor [F(1, 90) = 0.02, p = 0.89]. There was a significant CFA x THC interaction effect [F(1, 90) = 4.01, p = 0.048] (**Fig. 7I**) in females, with an effect size of 0.045. Tukey’s post-hoc comparisons showed no significant pairwise differences. A 2-way ANOVA for R_input_ data collected from Delayed neurons showed no main effect of CFA [F(1,80) = 0.43, p = 0.51] or vapor [F(1, 80) = 0.06, p = 0.81], and no CFA x THC interaction effect [F(1, 80) = 1.52, p = 0.22] (**Fig. 7J**) in females.

In females, a 2-way ANOVA for SAG data collected from Onset neurons revealed no main effect of CFA [F(1,93) = 0.01, p = 0.93] or vapor [F(1, 93) = 1.35, p = 0.25], and no CFA x THC interaction effect [F(1, 93) = 0.4, p = 0.53] (**Fig. 7K**). A 2-way ANOVA for SAG data collected from Delayed neurons revealed no main effect of CFA [F(1, 79) = 2.64, p = 0.11] or vapor [F(1,79) = 0.22, p = 0.64], and no CFA x THC interaction effect [F(1, 79) = 0.13, p = 0.72] (**Fig. 7L**) in females.

In males, firing rate response curves from Onset neurons of different experimental groups appeared to overlap (**Fig. 8A**). A 2-way ANOVA for firing rate @ 150 pA of Onset neurons revealed no main effect of CFA [F(1,115) = 0.08, p = 0.78] or vapor [F(1,115) = 0.31, p = 0.58], and no CFA x THC interaction effect [F(1,115) = 0.78, p = 0.38] (**Fig. 8B**) in males. A 2-way ANOVA for firing rate gain in Onset neurons revealed no main effect of CFA [F(1,115) = 0.1, p = 0.75] or vapor [F(1,115) = 0.01, p = 0.91], and no CFA x THC interaction effect [F(1,115) = 0.89, p = 0.35] (**Fig. 8C**) in males.

**Figure 8:**
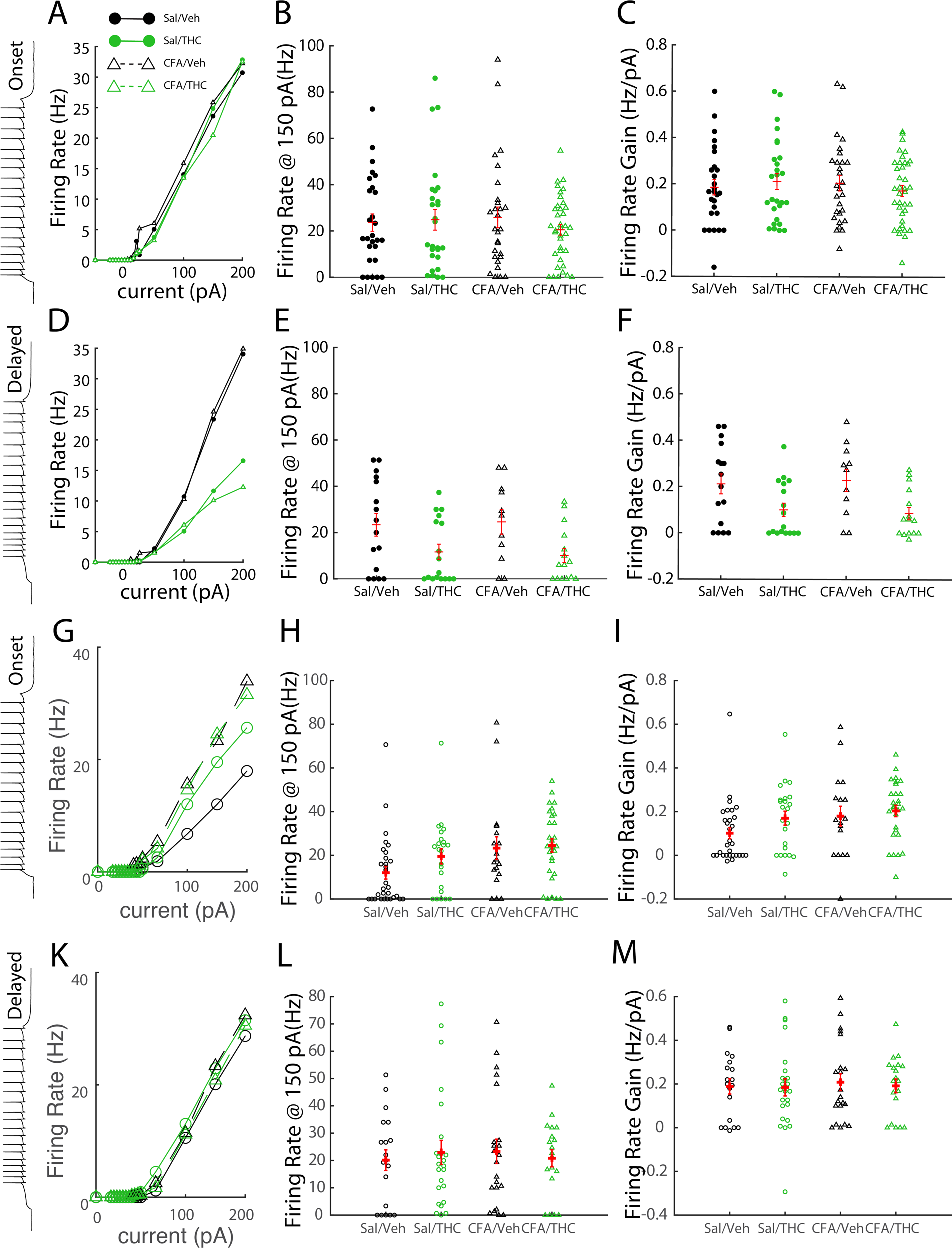
Firing phenotype-specific effects of CFA and vapor on action potential response properties. (**A**) Mean firing rate response curves of Onset vlPAG neurons from male rats to injected current. 2-Way ANOVAs show: (**B**) no effect of CFA [F(1,115) = 0.08, p = 0.7756] or vapor [F(1,115) = 0.31, p = 0.5814] on firing rate, and (**C**) no effect of CFA [F(1,115) = 0.1, p = 0.7546] or vapor [F(1,115) = 0.01, p = 0.9086] on firing rate gain in vlPAG Onset neurons. (**D**) Mean firing rate response curves of Delayed vlPAG neurons to injected current. (**E**) There was a main effect of vapor on firing rate in vlPAG Delayed neurons [F(1,55) = 9.78, p = 0.0028]. (**F**) There was a main effect of vapor on firing rate gain in vlPAG Delayed neurons [F(1,55) = 11.83, p = 0.0011]. (**G**) Mean firing rate response curves of Onset vlPAG neurons from female rats to injected current. 2-Way ANOVAs show: (**H**) a main effect of CFA [F(1,95) = 5.22, p = 0.0245] on firing rate. There is no effect of vapor [F(1,95) = 1.51, p = 0.2229] and no CFA x vapor interaction effect [F(1,95) = 0.79, p = 0.3768] on firing rate in neurons from female rats. (**I**) There is no effect of CFA [F(1,95) = 3.18, p = 0.0779] or vapor [F(1,95) = 2.16, p = 0.1449] on firing rate gain in vlPAG Onset neurons. (**K**) Mean firing rate response curves of Delayed vlPAG neurons to injected current. (**L**) There was no main effect of CFA [F(1,80) = 0.02, p = 0.895] or vapor [F(1,80) = 0.002, p = 0.9698] on firing rate in vlPAG Delayed neurons. (**M**) There was no main effect of CFA [F(1,80) = 0.02, p = 0.895] or vapor [F(1,80) = 0.002, p = 0.9698] on firing rate in vlPAG Delayed neurons from female rats.

In males, firing rate response curves from Delayed neurons from THC vapor-treated rats appeared to be lower than the curves from PG vehicle vapor-treated rats (**Fig. 8D**). A 2-way ANOVA for firing rate @ 150 pA of Delayed neurons revealed a main effect of vapor [F(1,55) = 9.78, p = 0.0028], with lower firing rates in the THC groups and an effect size of 0.151. There was no main effect of CFA [F(1,55) = 0.002, p = 0.97] and no CFA x THC interaction effect [F(1,55) = 0.11, p = 0.74] (**Fig. 8E**) in males. Tukey’s post-hoc pairwise comparisons revealed no significant differences between individual groups. A 2-way ANOVA for firing rate gain of Delayed neurons revealed a main effect of vapor [F(1,55) = 11.83, p = 0.0011], with lower firing rate gain in the THC groups and an effect size of 0.177. There was no main effect of CFA [F(1,55) = 9e^-4^, p = 0.98] and no CFA x THC interaction effect [F(1,55) = 0.18, p = 0.67] (**Fig. 8F**) in males. Post-hoc pairwise comparisons revealed no significant differences between individual groups.

In females, firing rate response curves from Onset neurons of different experimental groups appeared to overlap (**Fig. 8G**). A 2-way ANOVA analysis of firing rate @ 150 pA of Onset neurons revealed a significant main effect of CFA [F(1,95) = 5.22, p = 0.025] in females, with an effect size of 0.055. There was no main effect of vapor [F(1, 95) = 1.51, p = 0.3768], and no CFA x THC interaction effect [F(1, 95) = 0.79, p = 0.38] (**Fig. 8H**) in females. Tukey’s post-hoc pairwise comparisons revealed a significant difference between Sal/Veh and CFA/Veh groups (p < 0.05). A 2-way ANOVA for firing rate gain in Onset neurons revealed no main effect of CFA [F(1, 95) = 3.18, p = 0.078] or vapor [F(1, 95) = 2.16, p = 0.14], and no CFA x THC interaction effect [F(1, 95) = 0.52, p = 0.47] (**Fig. 8I**) in females. A 2-way ANOVA for firing rate @ 150 pA of Delayed neurons showed no main effect of CFA [F(1,80) = 0.02, p = 0.90], no main effect of vapor [F(1, 80) = 0.002, p = 0.97] and no CFA x THC interaction effect [F(1, 80) = 0.42, p = 0.52] (**Fig. 8K,L**) in females. A 2-way ANOVA for firing rate gain of Delayed neurons revealed no main effect of CFA [F(1, 80) =0.09, p = 0.77], no main effect of vapor [F(1,80) = 0.12, p = 0.73] and no CFA x THC interaction effect [F(1, 80) = 0.03, p = 0.87] (**Fig. 8M**) in females.

#### CFA and THC effects on synaptic properties in vlPAG Onset and Delayed neurons

We applied the same phenotype classification to the synaptic data to determine if different type of neurons exhibited different effects of CFA and vapor on synaptic properties in vlPAG neurons. In males, a 2-way ANOVA for sIPSC amplitude data collected from Onset neurons revealed no main effect of CFA [F(1,101) = 1.13, p = 0.29] or vapor [F(1,101) = 2.3, p = 0.13], and no CFA x THC interaction effect [F(1,101) = 0.002, p = 0.97] (**Fig. 9A**) in males. A 2-way ANOVA analysis of sIPSC amplitude data collected from Delayed neurons revealed a main effect of vapor [F(1,45) = 16.99, p = 0.0002] with lower sIPSC amplitudes in the THC groups and an effect size of 0.274. There was no effect of CFA [F(1,45) = 1.28, p = 0.26] and no CFA x THC interaction effect [F(1,45) = 0.17, p = 0.68] (**Fig. 9B**) in males. Tukey’s post-hoc pairwise comparisons revealed significant differences between Sal/Veh and Sal/THC groups (p < 0.01) and Sal/Veh and CFA/THC groups (p < 0.01).

**Figure 9:**
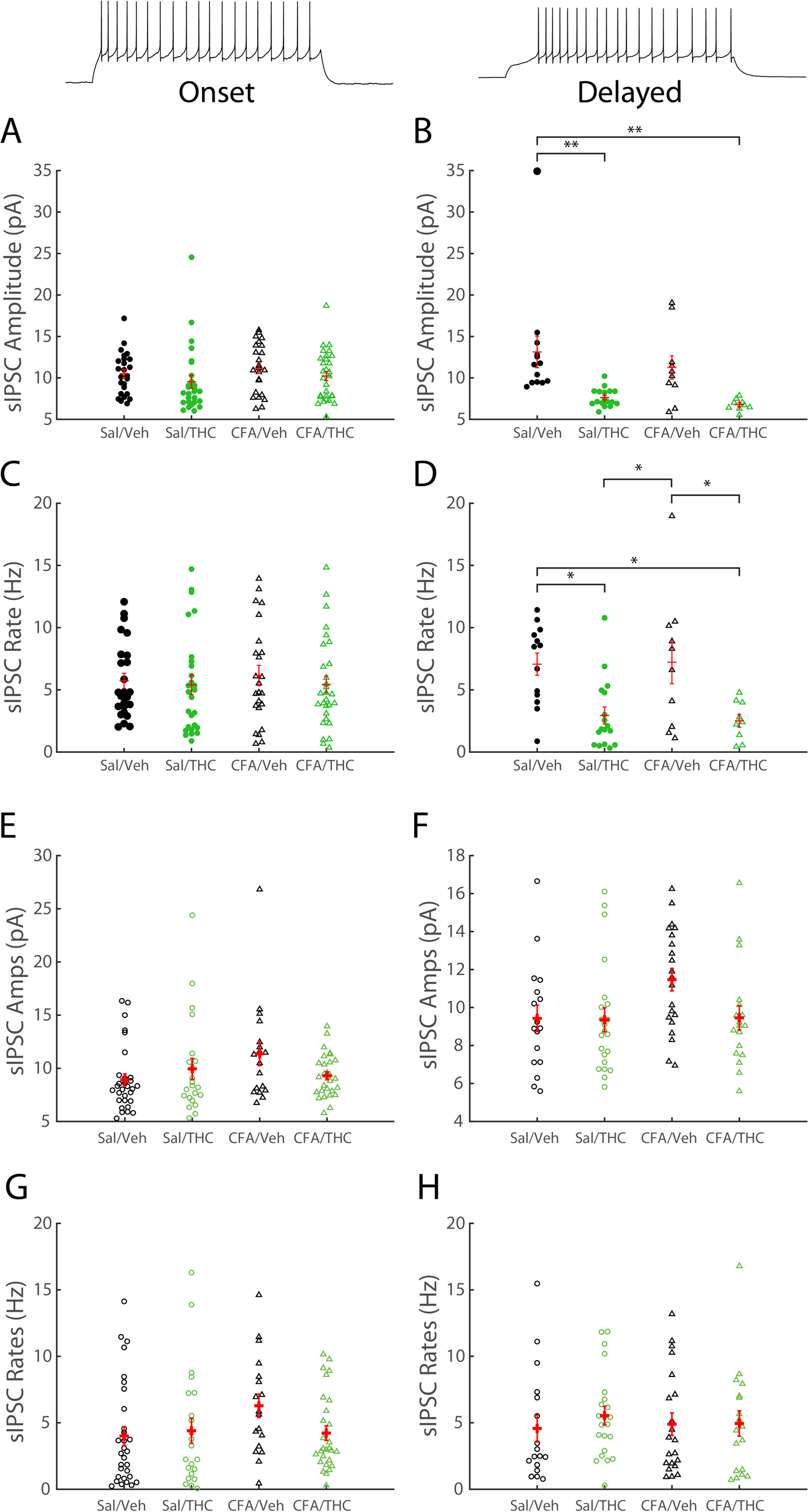
Firing phenotype-specific effects of CFA and vapor on synaptic properties of vlPAG neurons. (**A**) 2-Way ANOVA shows no main effect of CFA [F(1,101) = 1.13, p = 0.291] or vapor [F(1,101) = 0.002, p = 0.9664] on sIPSC amplitude in Onset vlPAG neurons from male rats. (**B**) There was a main effect of vapor on sIPSC amplitude in vlPAG Delayed neurons [F(1,45) = 16.99, p = 0.0002]. (**C**) There was no main effect of CFA [F(1,101) = 0.1, p = 0.7531] or vapor [F(1,101) = 0.51, p = 0.4761] on sIPSC rate in vlPAG Onset neurons. (**D**) There was a main effect of vapor on sIPSC rate in vlPAG Delayed neurons [F(1,45) = 18.8, p = 0.0001]. (**E**) 2-Way ANOVA shows no main effect of CFA [F(1,94) = 1.46, p = 0.2296] or vapor [F(1,94) = 0.5, p = 0.4805] on Sipsc amplitude in Onset vlPAG neurons from female rats. There is a CFA x vapor interaction effect [F(1,94) = 438, p = 0.0391] on sIPSC amplitude in vlPAG neurons from female rats. (**F**) There is no main effect of CFA [F(1,74) = 2.77, p = 0.1004] or vapor [F(1,74) = 2.63, p = 0.1008] on sIPSC amplitude in vlPAG Delayed neurons from female rats. (**G**) There was no main effect of CFA [F(1,94) = 1.84, p = 0.1786] or vapor [F(1,94) = 1.16, p = 0.2837] on sIPSC rates in vlPAG Onset neurons. (**I**) There was no main effect of CFA [F(1,74) = 0.03, p = 0.8743] or vapor [F(1,74) = 0.35, p = 0.5584] on sIPSC rates in vlPAG Delayed neurons from female rats. *p < 0.05, **p < 0.01.

In males, a 2-way ANOVA for sIPSC rate data collected from Onset neurons revealed no main effect of CFA [F(1,101) = 0.1, p = 0.75] or vapor [F(1,101) = 0.51, p = 0.48] and no CFA x THC interaction effect [F(1,101) = 0.07, p = 0.79] (**Fig. 9C**) in males. A 2-way ANOVA for sIPSC rate data collected from Delayed neurons revealed a main effect of vapor [F(1,45) = 18.8, p = 0.0001] with lower sIPSC rates in the THC groups and an effect size of 0.295. There was no effect of CFA [F(1,45) = 0.02, p = 0.90] and no CFA x THC interaction effect [F(1,45) = 0.08, p = 0.78] (**Fig. 9D**) in males. Tukey’s post-hoc pairwise comparisons revealed significant differences between Sal/Veh and Sal/THC groups (p < 0.05), Sal/Veh and CFA/THC groups (p < 0.05), Sal/THC and CFA/Veh groups (p < 0.05) and CFA/Veh and CFA/THC groups (p < 0.05).

In females, a 2-way ANOVA analysis of sIPSC amplitude data collected from Onset neurons revealed no main effect of CFA [F(1,97) = 1.46, p = 0.23] or vapor [F(1, 97) = 0.5, p = 0.48]. There was a significant CFA x THC interaction effect [F(1, 97) = 4.38, p = 0.039] (**Fig. 9D**) in females, with an effect size of 0.046. Tukey’s post-hoc pairwise comparisons showed no significant pairwise differences between group means. A 2-way ANOVA for sIPSC amplitude data collected from Delayed neurons showed no significant main effect of vapor [F(1,77) = 2.77, p = 0.10] or CFA [F(1, 77) = 2.63, p = 0.11] and no CFA x THC interaction effect [F(1, 77) = 2.28, p = 0.14] (**Fig. 9E**) in females.

In females, a 2-way ANOVA analysis of sIPSC rate data collected from Onset neurons showed no main effect of CFA [F(1,94) = 1.84, p = 0.18] or vapor [F(1, 94) = 1.16, p = 0.28] and no CFA x THC interaction effect [F(1, 94) = 2.52, p = 0.12] (**Fig. 9G**). A 2-way ANOVA analysis of sIPSC rate data collected from Delayed neurons showed no main effect of vapor [F(1,74) = 0.03, p = 0.87], no effect of CFA [F(1,74) = 0.35, p = 0.5584] and no CFA x THC interaction effect [F(1, 74) = 0.28, p = 0.60] (**Fig. 9H**) in females.

## Discussion

Although cannabinoids have been identified as a possible alternative to opioids for the treatment of chronic pain, little is known about the site(s) and mechanism(s) of action for THC vapor inhalation effects on pain-related outcomes, nor about THC vapor dosing regimens and/or duration of THC vapor effects on pain-related outcomes. This study was designed to examine the anti-nociceptive effects of THC exposure using an animal model of chronic inflammatory pain (CFA). The current study shows that chronic exposure to vaporized THC (100 mg/mL) one hour daily for 10 consecutive days acutely and repeatedly rescues CFA-induced thermal hyperalgesia in male and female Wistar rats, and that this effect lasts at least 24 hours. Chronic THC vapor inhalation also rescued mechanical hypersensitivity in CFA-treated male rats, but not in CFA-treated female rats, and this effect persisted in males 24 hours after the last THC vapor session. It is not clear why THC vapor inhalation did not reduce mechanical allodynia in CFA-treated female rats, but the THC-plasma levels reported here match the levels reported in a prior study by our group that showed substantial brain penetrance of vaporized THC and its metabolites in female (and male) rats (Baglot et al., 2021). There was a general reduction of mechanical sensitivity following both THC and vehicle vapor exposure in all females, regardless of CFA status – we did not observe this effect in thermal nociception assays, and we did not observe this effect in males. It is possible that vapor exposure or transportation to the testing environment produced some degree of stress-induced analgesia in female rats.

We report that chronic THC vapor exposure alters intrinsic and synaptic properties of vlPAG neurons in male rats, whereas CFA treatment did not affect these outcome measures. Specifically, THC vapor history reduced firing rate, firing rate gain, sIPSC amplitude and sIPSC frequency in vlPAG neurons of rats sacrificed 24 hours after termination of the 10^th^ day of vapor exposure. Furthermore, these effects were found specifically in vlPAG neurons that exhibited a delayed-firing phenotype, but not in neurons that exhibited an onset-firing phenotype. Quantification of effect sizes showed that the reduction of inhibitory synaptic amplitude and rates were approximately twice as large as reductions in the firing rates of the same neurons. This suggests that chronic THC vapor inhalation may activate descending pain modulation pathways via disinhibition of delayed-firing vlPAG neurons.

In female rats, we showed that CFA increases firing rates specifically in Onset-firing type neurons. We also show that CFA increases sIPSC amplitudes and that chronic THC rescues this increase and that this effect is likely driven by Onset-type neurons. Prior work also showed that CFA alters tonic GABAergic currents and miniature spontaneous postsynaptic current events in vlPAG neurons of female rats, but not male rats (Tonsfeldt et al., 2016). More recently, it was reported that CFA increases action potential firing rates in phasic-firing vlPAG neurons recorded from male and female rats (McPherson et al., 2021). Our study shows some similar results to these studies: specifically, CFA increase sIPSC amplitudes and firing rates in Onset (i.e. phasic) vlPAG neurons from female rats. In contrast with prior work, our recordings showed no firing rate changes in vlPAG neurons from CFA-treated male rats.

The CB1 receptor is highly expressed throughout the periaqueductal gray (Wilson-Poe et al., 2012). CFA treatment leads to a decrease in CB1R-mediated suppression of synaptic inhibition in vlPAG of rats (Bouchet and Ingram, 2020), and repeated cannabinoid exposures may downregulate CB1R in the rat brain (Breivogel et al., 1999). Our observation of chronic THC vapor-induced suppression of spontaneous synaptic inhibition in vlPAG (independent of pain condition) may suggest one potential mechanism whereby cannabinoids exert opioid-sparing effects in male rats – specifically chronic THC vapor may disinhibit RVM-projecting neurons in vlPAG.

As discussed above, prior work using THC injections reported that systemic and intra-plantar THC injection reduces thermal hyperalgesia and/or mechanical allodynia in CFA-treated rats, and these effects are stronger in female rats (Craft et al., 2013). Other studies from the same group reported that systemically injected THC reduces pain-related behaviors in CFA-treated male and female rats but does not reduce hindpaw edema, with little evidence for sex differences or tolerance to those effects after repeated treatment (Britch et al., 2020; Britch and Craft, 2023). One recent study reported that vaporized THC-dominant cannabis extract reduces mechanical allodynia and paw edema similarly in male and female rats treated with CFA, and that tolerance to these effects develop with chronic exposure (Craft et al., 2023). Here, we reported anti-allodynic and anti-hyperalgesic effects of THC vapor inhalation in male rats, but only anti-hyperalgesic effects of THC vapor inhalation in female rats, and we did not observe tolerance to the anti-nociceptive effects of chronic THC vapor inhalation. There are some important methodological differences between these studies that may account for the discrepant patterns of results across studies: first, we used Wistar rats whereas prior studies all used Sprague-Dawley rats; second, we tested the effects of vaporized THC whereas prior studies tested the effects of systemically injected and intraplantar injected THC, as well as the effects of vaporized cannabis extract; third, chronic treatment in this study constituted 10 days of THC vapor exposure whereas prior studies administered THC injections or vaporized cannabis extracts for 3-4 days; finally, we injected the hindpaw of rats with 150 uL of CFA whereas prior studies injected the hindpaw rats with 100 uL of CFA.

Previous studies reported anti-nociceptive effects of vaporized THC in “pain-naïve” rats using the tail flick assay at this time point (Javadi-Paydar et al., 2018; Nguyen et al., 2018); we did not replicate this effect, but we also used different behavioral assays (Hargreaves and von Frey). One recent study from the same group reported that maximal anti-nociceptive effects occurred 60 minutes after the end of THC vapor exposure in male Wistar rats (Moore et al., 2021). Aside from the fact that we used different nociception assays in the present set of studies, it is possible that observed THC vapor effects may have changed if we delayed testing for thermal nociception and mechanical sensitivity. It is important to re-emphasize that THC vapor did attenuate thermal hyperalgesia in CFA-treated male and female rats, and attenuated mechanical hypersensitivity in CFA-treated male rats, but not CFA-treated female rats. Interestingly, a recent paper reported an anti-allodynic effect of systemically injected THC in CFA-treated female (and male) rats (Britch and Craft 2021), which suggests that route of THC administration may be important for mediating the effect of THC on mechanical allodynia in CFA-treated female rats. There are also differences in rat strain, and in the time course of drug administration and behavioral testing in these two studies that may account for these discrepant results.

In the periphery, different nociceptors are responsible for transmitting different types of pain information. For example, selective ablation of nociceptors expressing Mas-related G protein-couple receptor subtype D (MRGPRD) attenuates mechanical hypersensitivity (Cavanaugh et al., 2009), while ablation of nociceptors expressing calcitonin gene-related peptide (CGRP) attenuates thermal nociception (McCoy et al., 2013). More recent evidence suggests that specific cells and circuits in the CNS have distinct roles in mediating different pain modalities (Carrasquillo & Gereau, 2007; Singh et al., 2022; Wilson et al., 2019. For example, chemogenetic activation of all vlPAG neurons, inhibition of vlPAG GABA neurons and activation of vlPAG glutamate neurons each modulate thermal nociception but not mechanical sensitivity in mice (Samineni et al., 2017). Future work will identify the specific cells and circuits that are responsible for mediating the modality-specific rescue effects of THC vapor inhalation in female rats treated with CFA.

The specific contributions of GABAergic and glutamatergic synaptic transmission to antinociception mediated by descending vlPAG projections to RVM are not clear. Anatomical studies have reported that a majority (∼71%) of vlPAG fibers that contact RVM neurons are GABAergic, and that this proportion is observed in ON cells, OFF cells and neutral cells in the RVM of mice (Morgan et al., 2008). In contrast, another study in mice reported that GABAergic neurons in the vlPAG were non-overlapping with retrogradely labeled RVM-projecting neurons, and that opioid-mediated analgesia depended on suppression of spiking activity of GABAergic neurons in the vlPAG, but that this effect was independent of presynaptic suppression of inhibition onto RVM-projecting neurons (Park et al., 2010). As mentioned above, chemogenetic activation of vlPAG glutamate neurons and chemogenetic activation of vlPAG GABA neurons each promote nociception in mice (Samineni et al., 2017). To our knowledge, there are currently no data from paired recordings that demonstrate a microcircuit involving a local inhibitory interneuron and an excitatory RVM-projecting neuron in the vlPAG.

Our study represents the first characterization of late-firing neurons in the vlPAG of rodents. In the BLA, late-firing neurons comprise ∼22% of parvalbumin-positive (PV^+^) local inhibitory interneurons (Woodruff and Sah, 2007). Late-firing phenotype neurons have been observed in the central amygdala, primarily in the lateral subdivision (Martina et al., 1999; Lopez de Armentia and Sah, 2004; Amano et al., 2012). A subset of both protein kinase C delta-positive (PKCδ^+^) and somatostatin-positive (SOM^+^) neurons in the CeA of mice exhibit late-firing phenotypes (Ciocchi et al., 2010; Haubensak et al., 2010; Amano et al., 2012). In the CeA of mice, SOM^+^ late-firing neurons have higher excitability than PKCδ^+^ late-firing neurons, but this difference is abolished in a sciatic nerve cuff model of neuropathic pain 6-14 days after injury (Adke et al., 2021). In the same model of neuropathic pain, PKCδ^+^ CeA neurons were pro-nociceptive and this seemed to be due to an increase in excitability of late-firing PKCδ^+^ but not regular-spiking PKCδ^+^ neurons (Wilson et al., 2019). Finally, in a mouse model of chronic inflammatory pain, late-firing vlPAG-projecting neurons in the CeA exhibited no changes 1 day after CFA injection into the hindpaw (Li and Sheets, 2018). It is not clear how cell firing phenotypes of different vlPAG neuron subsets is related to 1) projection targets of those neurons, 2) alterations by chronic inflammatory (or neuropathic) pain, 3) THC sensitivity, and 4) pain-related behaviors, or combinations of these factors. For example, if the role of these cells is similar to that of late-firing PKCδ^+^ neurons in the CeA (i.e., pro-nociceptive), then reduction of their firing activity via reductions in firing rate response and firing rate gain may have anti-nociceptive effects.

Here, we showed that CFA produces thermal hyperalgesia and mechanical hypersensitivity in adult male and female Wistar rats. Chronic THC vapor inhalation reversed thermal hyperalgesia in male and female rats treated with CFA, rats but reversed mechanical hypersensitivity only in male rats treated with CFA. Chronic THC vapor inhalation also modulated intrinsic and synaptic properties of vlPAG neurons in male rats, and some of these effects were specific to neurons with a delayed-firing phenotype. CFA increased presynaptic inhibition and firing rates in Onset neurons from female rats, and THC rescued the CFA-induced increase in firing rates in these neurons. Future work will determine the potential role of specific vlPAG cells and circuits in mediating chronic inflammation and THC vapor inhalation effects on pain-related outcomes.

## Funding

This work was supported in part by a Merit Review Award #I01 BX003451 (NWG) from the United States (U.S.) Department of Veterans Affairs, Biomedical Laboratory Research and Development Service, and by an NIH grant AA023305 (NWG).

## Conflict of interest statement

Dr. Gilpin owns shares in Glauser Life Sciences, Inc., a company with interest in developing therapeutics for mental health disorders. There was no direct link between those interests and the work contained herein.

## Acknowledgements

All Δ^9^-tetrahydrocannabinol (THC) was supplied in ethanolic solution (200mg/mL) by the U.S. National Institute on Drug Abuse (NIDA) Drug Supply Program.

## Notes

### Competing Interest Statement

The authors have declared no competing interest.

### Summary of Updates

The manuscript was expanded from the prior version to include all of the previous assays on females as well as males.

